# Structural basis for continuous DNA-end protection during ligation of double-strand breaks in yeast Non-Homologous End-Joining

**DOI:** 10.64898/2026.03.12.711350

**Authors:** Sophia Missoury, Claire Tettaravou, Sara Castelli, Amandine Pelletier, Vincent Morin, Paloma Fernández Varela, Virginie Ropars, Stefano Mattarocci, Pierre Legrand, Mauro Modesti, Stéphane Marcand, Jean-Baptiste Charbonnier, Marc Delarue

## Abstract

Non-homologous end joining (NHEJ) repairs DNA double-strand breaks by synapsing and ligating DNA ends. In vertebrates, DNA-PKcs promotes end alignment and processing, yet several eukaryotes - including budding yeast - perform NHEJ without DNA-PKcs, and the underlying mechanism remains unclear. Here we report cryo-electron microscopy structures of reconstituted *Saccharomyces cerevisiae* NHEJ synaptic complexes assembled on DNA ends bearing either 4-bp terminal microhomologies or blunt termini. On the microhomology substrate with a single 5’-P, we capture a ligation-competent short-range complex in which a single Dnl4 catalytic core is fully ordered and engaged at the 5′-adenylated nick. When two 5’-P are present, we resolve two predominant, coexisting DNA-aligned protective states in which both Dnl4 DBD-NTD modules occupy the break region with reciprocal geometries, supporting an alternating engagement model for sequential sealing of the two strands while maintaining continuous end association. In contrast, blunt-ended substrates yield a non-aligned protective configuration in which two Dnl4 catalytic modules constrain the DNA ends ∼30 Å apart in a ligation-incompatible arrangement, providing a structural explanation for slow blunt-end joining in yeast. Together, these structures define architectural and mechanistic principles of DNA-PKcs-independent NHEJ.

## Introduction

DNA double-strand breaks (DSBs) represent one of the most cytotoxic forms of DNA damage, threatening genome integrity and cell viability^1^. They arise from endogenous sources such as replication fork collapse, reactive oxygen species, or from exogenous attacks such as ionizing radiation and genotoxic chemicals^2–5^. Eukaryotic cells repair DSBs primarily through two conserved pathways: homologous recombination (HR) and non-homologous end joining (NHEJ)^6–8^. HR uses a homologous duplex as template, typically a sister chromatid, and therefore provides a slow, post-replicative mode of repair, whereas NHEJ directly rejoins DNA ends without requiring a template.

In eukaryotes, NHEJ is initiated by the recruitment of the Ku70/Ku80 (Ku) heterodimer to DNA ends, serving as a platform for the assembly of downstream repair factors^9–12^. Beyond this conserved initiation step, the subsequent stages of NHEJ - which include DNA end processing by nucleases or polymerases and ligation - diverge between species^13^. In vertebrates, Ku recruits the DNA-dependent protein kinase catalytic subunit (DNA-PKcs), forming the DNA-PK holoenzyme, and recruits Ligase IV (Lig4) and XRCC4 (as a subcomplex L4X4) as well as XLF to form a Long-Range (LR) synapsis complex that maintains the two DNA ends separate but in proximity^14–18^. Autophosphorylation of DNA-PKcs orchestrates a cascade of events through its serine/threonine kinase activity^19,20^, facilitates DNA end processing and promotes access to specialized enzymes such as the nuclease Artemis^21^ and possibly polymerase µ in a template-free activity^22,23^. These events can lead to the finding of a short micro-homology (MH) region between the DNA ends, ultimately enabling the transition to a short-range (SR), ligation-competent complex composed of Ku, XRCC4-Ligase IV, and XLF. Structural studies of this complex in the human system have revealed a dynamic multimeric architecture resembling an ω-shaped arrangement (or W-shaped), that stabilizes DNA ends and can accommodate diverse end structures during ligation^15–17,24–29^. Additional factors, including PNKP and the DNA polymerases μ and λ, contribute to end processing ^22,30–35^, while PAXX enhances NHEJ complex stability ^29,36^.

However, NHEJ can proceed relatively efficiently in mammalian cells lacking DNA-PKcs when DNA ends require minimal processing, indicating that DNA-PKcs is not an essential component of the core ligation machinery ^37,38^. The question arises as to how NHEJ proceeds in this case, at the structural level. On the evolutionary level it would be interesting to find out how a primitive version of the NHEJ system works without some of its co-factors.

In contrast to vertebrates, the budding yeast *Saccharomyces cerevisiae* lacks DNA-PKcs and the accessory NHEJ factor PAXX. Instead, NHEJ relies on the core machinery made of Ku (Yku70/Yku80 heterodimer)^39^, DNA ligase IV (Dnl4), its XRCC4-like partner Lif1, and the XLF-like factor Nej1 ^40–46^. In addition, the Mre11-Rad50-Xrs2^NBS1^ DNA repair complex (MRX in *S. cerevisiae*, MRN in other eukaryotes) contributes to Ku-dependent NHEJ in yeast ^47^ by promoting DNA ends tethering, while Xrs2 further stimulates DNA ligation through an interaction with Lif1, similar to the reported interaction between NBS1 and XRCC4 in human ^48,49^.

Despite these adaptations, budding yeast NHEJ (yNHEJ) is inefficient or slow at repairing blunt-ended DSBs *in vivo* ^50,51^. This limitation suggests intrinsic structural constraints within this DNA-PKcs-independent end-joining machinery.

While vertebrate DNA-PKcs-dependent NHEJ has been extensively characterized through biochemical and structural studies, the mechanistic details of DNA-PKcs-independent systems remain comparatively underexplored. However, the relative simplicity of genetic tools in yeast makes it an attractive target to dissect the yNHEJ mechanism through structural, biochemical and *in vivo* approaches.

Here, we reconstitute the core yeast NHEJ ligation machinery and establish its ability to repair DSBs with distinct end configurations. Using cryo-electron microscopy, we determine structures of yeast NHEJ ligation complexes bound to DNA ends, including blunt-ended DSBs at resolutions ranging from 3.3 to 5.8 Å. These structures reveal the architectural principles governing DNA-PKcs-independent end joining and show that Dnl4 forms intermediate protective synaptic complexes, using both catalytic cores to protect DNA ends prior to the recruitment of processing enzymes such as nucleases or polymerases until sequential ligation, structurally compensating for the absence of DNA-PKcs. Complementary genetic analyses validate key protein - protein interfaces identified by our structural studies.

## Results

### Structure of the ligation-competent synaptic complex in yeast NHEJ

Yeast NHEJ DNA ligase Dnl4 is a 941-amino-acids ATP-dependent ligase that catalyses the phosphodiester bond formation between the two DNA strands via the adenylation of the 5′ phosphate terminus on one side and its subsequent attack by the 3’-OH group on the other side. Dnl4 is made of a catalytic module containing three domains and a C-terminal region that contains two BRCT domains (**Fig. 1a**), that form a complex with Lif1 (XRCC4 in vertebrates). The catalytic module contains a DNA-binding domain (DBD), the nucleotidyltransferase domain (NTD) which performs the first adenylation reaction, and an OB-fold domain (OBD) that locks the ligase in an active conformation required for AMP transfer to the DNA end. Yeast Dnl4 and human Ligase IV share 27% sequence identity (**Extended Data Fig. 1**). While the DBD domain of one of the ligases is essential for the stability of the short-range (SR) synapsis complex in human ^24^, its NTD and OBD are only ordered with compatible DNA substrates, while the second catalytic core is usually not ordered. At the sequence level yeast Dnl4 from the *Saccharomyces* clade (*Z. fermentati*, *K. lactis*, *C. glabrata*) have three major segments that differ in human and vertebrate Ligase IV called Insert1, Insert2 and Insert3, whose role will become apparent in this study but can be placed already in a structural context using an Alphafold model of Dnl4 (**Extended Data Fig. 1a,c**). *S. cerevisiae* Yku80 and Yku70 lack the C-terminal and SAP domains respectively, which are critical for recruiting DNA-PKcs and stabilizing chromatin interactions (**Fig. 1a**). In addition, the yeast XLF homologue Nej1 lacks the mammalian canonical Ku-binding motif (KBM), which is responsible for its recruitment by Ku and stabilizes an open conformation of the vWA domain of Ku80 (**Extended Data Fig. 7**).

**Fig. 1.**
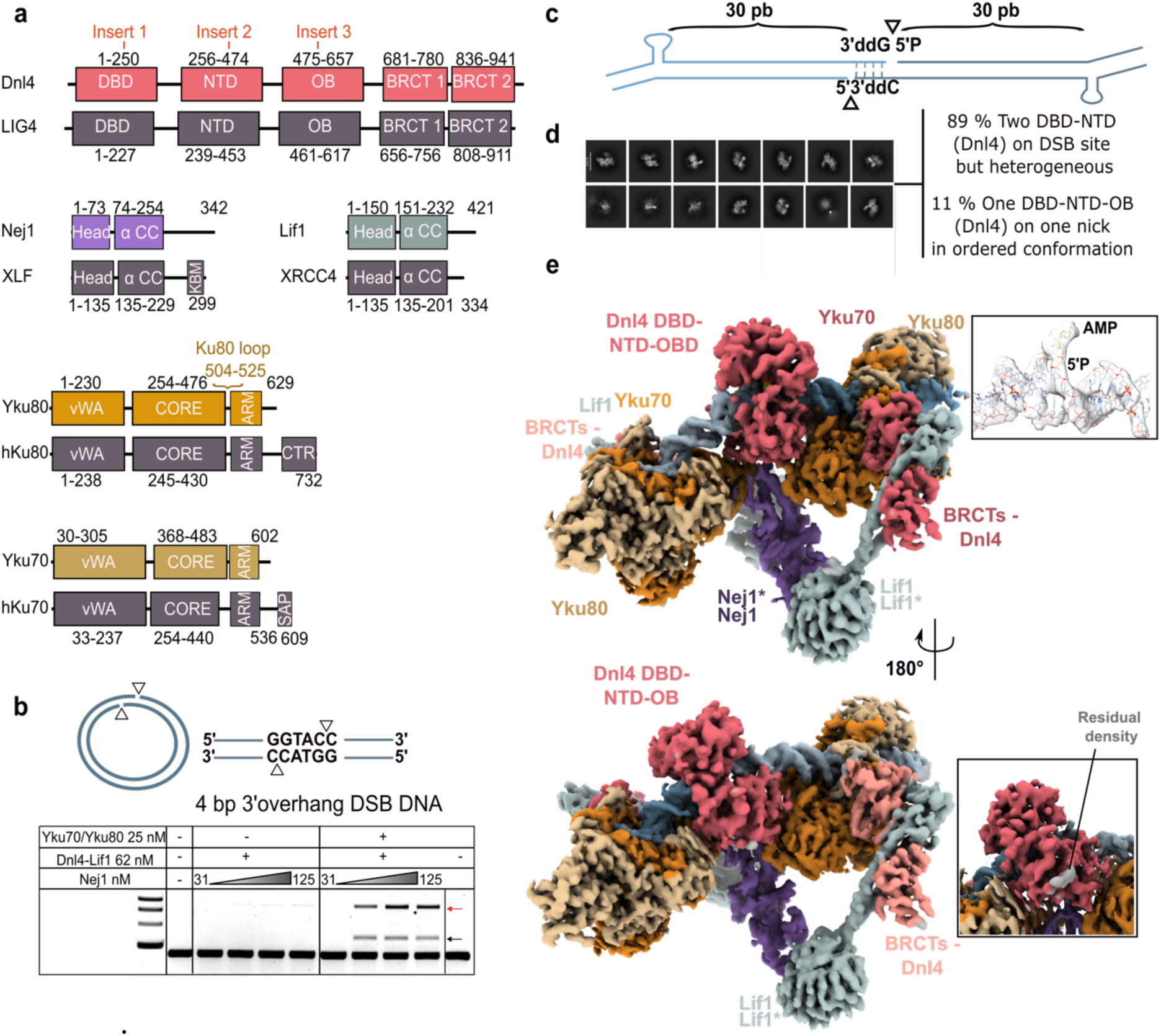
Architecture of the ligation-competent yeast end-joining synaptic complex. **a**, Domain organization of core NHEJ factors in yeast and human. The synaptic complex includes two molecules of Dnl4, one Nej1 homodimer, two Lif1 homodimers, and two Ku70/Ku80 heterodimers, which together stabilize double-strand break (DSB) DNA substrates. The loops or insertions in the DBD (Insert1), NTD (Insert2 and Insert3) of Dnl4 and in Ku80 (Loop) discussed in the text are indicated in colour. **b,** In vitro ligation assays performed using cohesive-ended DNA substrates (linear pUC19 2,686 bp) processed by yeast NHEJ (yNHEJ) factors at the indicated concentrations. Nej1 concentrations were titrated evenly from 31 to 125 nM. All tested NHEJ factors are required for efficient ligation: only a small amount of DNA is ligated in the absence of Ku and Nej1 is required in a concentration-dependent manner for DNA ligation of both cohesive-ended and blunt-ended substrates. The size markers come from the 1 kb NEB ladder (the lowest band is at 3 kb, then 4, 5, 6 kb). Intra and inter ligation products are indicated by a black-arrow and a red-arrow respectively. **c**, Schematic representation of the DNA substrate used to reconstitute the yeast NHEJ synapsis complex by cryo-EM. The substrate consists of two 30-nucleotide duplexes forming a cohesive-end DSB, one strand bearing a 5′ phosphate (5’-P) upstream and a 3′ dideoxyguanine (3’-ddG) modification downstream to prevent the ligation reaction to take place. The second strand has a 5’-OH upstream and a 3’-dideoxycytidine (3’-ddC) downstream. **d**, Representative cryo-EM 2D class averages of yNHEJ factors assembled with the DNA substrate shown in **c**. Out of 10 3D classes, one of them shows a completely ordered and active Dnl4 conformation, whose composite cryo-EM map is shown in **e**, color-coded as in **a**. The remaining 9 classes show two DBD-NTD present in cryo-EM density but with varying positions with respect to DNA ends, not easily distinguishable. They are representative of an intrinsic dynamic behaviour at the synapsis. **e**, Composite map of the best ordered conformation poised to react on the first nick. Each DNA duplex is bound by a Ku heterodimer (Yku70, orange; Yku80, light orange). Ku interacts with the BRCT1 and BRCT2 domains of Dnl4 (dark pink), which associates with Lif1 homodimers (grey-green). The Nej1 homodimer (purple) bridges two Lif1 homodimers through head-to-head interactions, stabilizing the centre of the short-range synaptic complex. The DBD-NTD-OBD module of one Dnl4 molecule is fully ordered and positioned for catalysis at one DNA end. Only the BRCT domains are seen in density for the second Dnl4 molecule. First inset, cryo-EM map consistent with a 5′-adenylated 5’-phosphate at the DNA terminus and a free K283 in Lig4. Second inset, cryo-EM density of an additional feature similar to the one seen in the human synaptic complex and assigned to the N-terminus of Ku70 (PDB ID 9N83).

To reconstitute the yNHEJ system, we purified *S. cerevisiae* Ku, Dnl4-Lif1 and Nej1, and determined the activity of each component using a plasmid-based ligation assay, involving 4-nt 3′ cohesive ends (KpnI-digested pUC19). Dnl4-Lif1 alone failed to ligate cohesive-end breaks after 60 minutes of incubation at 30 °C. The addition of Nej1 to the reaction led to detectable ligation products with a ratio of 2 Dnl4-(Lif1)_2_ to 1 (Nej1)_2_. Robust cohesive-end ligation required the Ku heterodimer (**Fig. 1b**).

To investigate by cryo-EM how Dnl4 operates on cohesive-compatible DNA ends, we used two 37-bp duplex substrates in which one strand contains a hairpin that orients Ku entry into DNA ends and prevents its translation (**Fig. 1c**). The substrates contain 3′-overhangs that are complementary in 4-bp forming two adjacent non-ligatable nicks (5′-P/3′-ddC and 5′-OH/3′-ddC), with a single 5′-phosphate available to isolate the first ligation event (**Fig. 1c**). Dnl4-Lif1, Ku, Nej1, and DNA were incubated with ATP and subjected to glycerol-gradient separation to assemble and stabilize the complex before vitrification (**Extended Data Fig. 2**). The cryo-EM map reveals that the general architecture of the synaptic complex closely resembles that of the human complex. However, we observe a continuum of DBD-NTD configurations in most particles (89%), with catalytic modules engaged at either the first nick, the second nick, or simultaneously at both nicks as a dimer (**Fig. 1d, Extended Data Fig. 2**). Nevertheless, ∼11% of particles display a single fully ordered Dnl4 catalytic module (DBD-NTD-OBD) bound to the DNA and position unambiguously the covalently linked AMP moiety at the 5′-phosphate, indicative of a ligation competent state (**Fig. 1d,e**). This observation is consistent with recent cryo-EM structures of the short-range human NHEJ complex ^17,24^ and with the crystallographic structure of human DNA-bound Lig4 ^52^. Cryo-EM reconstruction of the resulting active Dnl4 assembly yielded a *composite* map at an overall resolution of 3.3 Å (**Fig. 1e; Extended Data Fig. 2**). Local resolution analysis indicates a 3.0 Å resolution for the Ku-DNA region, whereas the Lif1-Nej1-BRCTs scaffold exhibited intermediate resolution (∼4.8 Å), indicative of a substantial conformational flexibility (**Extended Data Fig. 2 and Table 1**). Protein components corresponding to Yku80 (residues 1-588), Yku70 (22-569), Nej1 (1-254), Lif1 (1-236), the Dnl4 catalytic module (DBD-NTD-OBD; residues 1-653) and the BRCT domains of Dnl4 (residues 654-672) were unambiguously modelled, with unresolved regions corresponding primarily to solvent-exposed or intrinsically disordered segments. The correlation coefficients of the individual chains in the experimental map are indicated in **Extended Data Table 2**. An interesting region in human Lig4 is Insert1, located at residues 113-122 in the DBD, which distinguishes Lig4 from Lig1 and Lig3 and is well resolved in the electron density of the human ligation complex ^24^. In yeast Insert1 is shorter than in human and mediates a direct contact between Dnl4 (K121) and Nej1 (D177) and follows closely the AlphaFold prediction (**Extended Data Fig.1c**); it clearly differs from the human one and in the following we refer to the yeast version as type 2 and the human version as type 1. Insert1 can then be mapped to either type 1 or type 2 in other species intermediate between Saccharomycetales and vertebrates using AlphaFold models of Dnl4 and the structure of Lig4 bound to DNA (**Extended Data Fig.1b**). Insert3 is located next to Insert1 in space and its length is clearly anticorrelated to the one of Insert1.

Insert 2 is not completely resolved in this case but it clearly positions K362 near the second nick, consistent with a protective role as proposed by Liu *et al*. ^24^ and also with the AlphaFold prediction (**Extended data 1a**).

In the two following sections, we discuss in more details the architecture of this ligation-competent complex in which one Dnl4 molecule adopts an active conformation.

### Conserved Ku positioning in an open conformation in the yeast NHEJ synapsis complex

Two Ku heterodimers were positioned and refined in the cryo-EM map using the crystallographic structure of Ku alone as an initial model (PDB ID 5Y58). In the assembled yeast NHEJ complex, each Ku sits ∼15 bp away from the DNA break, resulting in a symmetric inter-dimer spacing of ∼258 Å (distance measured between two Yku80:D114) (**Fig. 2a**). This arrangement closely mirrors that of the human NHEJ SR synapsis (PDB ID 9N83 and 7LSY), where the two Ku molecules are separated by 260 Å. Consistent with what is observed in the human complex, Nej1 emerges as a central piece in the architectural scaffold of the yeast NHEJ machinery. Nej1 is a 342-amino acids homodimer that was manually fitted in the electron density map and refined without any major rearrangement (**Extended Data Fig. 2**). The central region of Nej1 (residues 144-184) adopts a long coiled-coil α-helix structure parallel to the equivalent helix of the second monomer that stabilizes the dimer (**Fig. 2a**). Residues 188-204 fold back towards the N-terminal head domain in three short α-helices oriented opposite to the central helix, as in human XLF, the functional counterpart of Nej1. The tip of Nej1 central α-helix makes a small contact with Yku70. Although this interface is modest in buried surface area (∼80 Å²), it contributes to the additional stabilization of the overall assembly through a direct hydrophobic contact between Nej1 L225 and Yku70 W186. The N-terminal domain of Nej1 adopts a globular structure, flanked by an anti-parallel β-sheet surrounded by an α-helix, which mediates key protein-protein interactions. The Lif1-Nej1-Lif1 assembly stabilizes DNA synapsis by tethering together the two Ku complexes on both sides of the DNA break (**Fig. 2a, b**). Each Ku forms a positively charged ring that encircles ∼20 bp of DNA, corresponding to roughly two helical turns (**Fig. 2c, d**). Protein-DNA interactions involve mainly the minor groove of the double-stranded DNA and implicate residues inside the Ku ring as well as the bridging β-turn-β loop encircling the DNA, as observed in human Ku bound to the DNA in the SR synapsis complex^53^ (**Fig. 2c**).

**Fig. 2.**
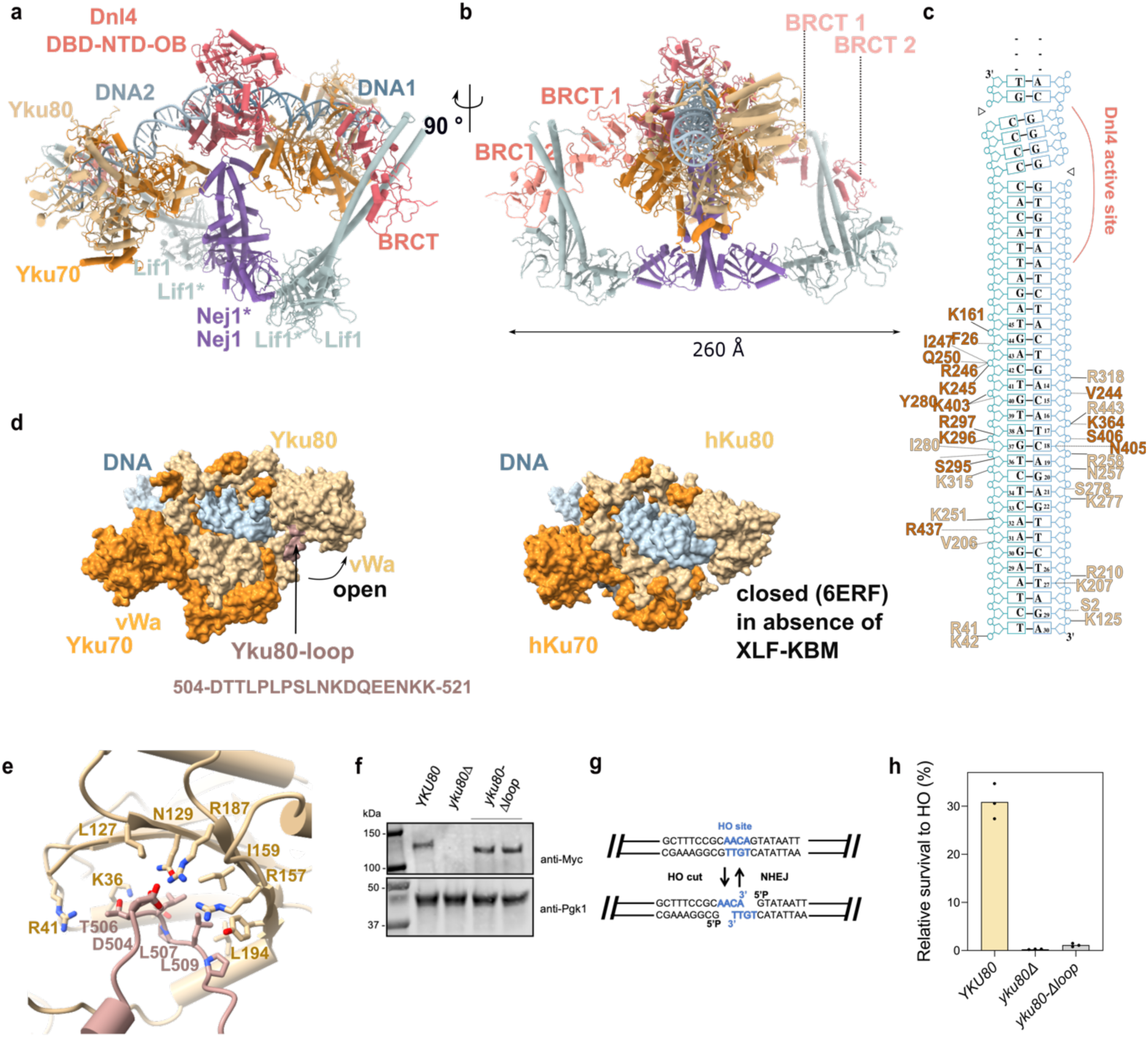
Protein-protein and protein-DNA interactions in Ku in the ligation-competent NHEJ synaptic complex. **a,** General architecture of the synaptic complex involved in the end-joining ligation assembly. Nej1 forms a homodimer (purple) that bridges two Lif1 homodimers (green) through head-to-head interactions. Each Lif1 dimer engages the BRCT1 and BRCT2 domains of Dnl4 (red) via its coiled-coil region, thereby tying the ligase to the rest of the complex. BRCT1 additionally contacts the Ku70/Ku80 heterodimer (red and light pink). Through its long coiled-coil and its orientation, the Nej1-Lif1 complex positions Ku70/Ku80 relative to the DNA-DSB. **b,** Front view of the Lif1-Nej1-Lif1 complex. Each Nej1 monomer binds to the adjacent Lif1 monomer, forming the arms of an extended μ-shaped architecture spanning approximately 260 Å in length. **c,** Detailed view of protein-DNA contacts between one Ku and one DNA duplex of the synapsis. The two nicks are indicated by open triangles. Key residues of Yku70 are shown in dark orange and for Yku80 in light orange: they interact primarily with the phosphate backbone, schematically shown as circles; some contacts are observed with the ribose moieties of the DNA and very few with bases. Hydrogens bonds are shown by a full line; non-bonded contacts are shown by dash-lines. **d,** Zoomed-in view on a single Yku complex that encircles 20 bp of the DNA duplex. Core domains of Yku stabilize the DNA duplexes (light blue). Despite the absence of a X-KBM motif in Nej1, the von Willebrand A (vWA) domain of Yku80 adopts an open conformation relative to the DNA-binding channel through the Yku80-loop (brown). For comparison at the right, the human KU (hKU) adopts a closed conformation for hKu80 in absence of XLF Ku-binding motif X-KBM (PDB ID 6ERF) **e,** Close-up view of the Yku80 pocket showing the interaction of Yku80 loop (in the range 504-513) which is not present in human Ku80 and occupies the position normally engaged by the X-KBM in the human complex. **f,** Western blot analysis of WT YKU80 and the *yku80-Δloop* mutant, both tagged with 13xMyc and detected using an anti-Myc antibody. **g,** Schematic representation of the NHEJ assay based on transient induction of the HO endonuclease. HO expression generates a site-specific DNA double-strand break, and cell survival is used as a readout of NHEJ efficiency. **h,** Relative survival following HO expression, normalised to the uninduced condition.

The C-terminal tail of Nej1 (residues 205-305) could not be modeled, consistent with predictions that this region is intrinsically disordered. In contrast to the human NHEJ assembly, we did not observe any density corresponding to a XLF-Ku binding motif (X-KBM) at the equivalent position on Yku80. In human cells this motif promotes an outward rotation of the Ku80 vWA domain, thereby converting Ku from a closed to an open conformation^54^. Sequence alignment of Nej1 with other XLF family members (including human XLF) indicates that the X-KBM motif is not conserved in yeast Nej1 (**Extended Data Fig. 7**). However, we mapped a well-defined internal loop in Yku80 from residues 505 to 525 in the cryo-EM density map that occupies the same position as the human X-KBM (**Fig. 2d, 2e**) seen in both the crystal structure of Ku (PDB ID 5Y58) and the cryo-EM structure of the Ku heterodimer in complex with double-stranded DNA (PDB ID 8S82) ^55^. This Yku80 loop may compensate for the absence of a Nej1-derived KBM in yeast, and structural superpositions indicate that it could stabilize the open conformation of the vWA domain (**Fig. 2d, 2e**). This open state has been suggested to play a role in the inward translocation of Ku along dsDNA^55^. To explore the functional contribution of the Yku80 loop, we assayed NHEJ efficiency in cells expressing a Yku80 mutant lacking this region (yku80-Δ510-523) (**Fig. 2f, 2g**). This mutation did not affect protein levels (**Fig. 2g**) but resulted in a severe defect in end-joining repair, phenocopying Yku80 absence (**Fig. 2h**). This result is consistent with the Yku80 loop being required for Ku function. In human cells, disruption of the Ku80 X-KBM pocket - which prevents stabilization of X-KBM and thereby blocks Ku80 opening, produces a similar severe defect, markedly increasing radiosensitivity ^54^. Yku80 loop is present in saccharomycetes clade but absent in *S. pombe*-like yeasts, confirming that the latter are more closely related to vertebrates than to sacchamorycetes clade (**Extended data Fig. 9**).

### Nej1-Lif1 assembly directs the DNA ends configuration during end-joining

Although the overall architecture of Nej1-Lif1-Dnl4_BRCTs_ complex mirrors the human XLF-XRCC4-Ligase IV_BRCTs_ complex, a marked divergence emerges in the overall direction of the Lif1 coiled-coil. Human XRCC4 undergoes a pronounced kink near residue L172 with a ∼20° bend in the local helical axis, resulting in an angle of ∼60° between XRCC4-XLF long helices relative to DNA (PDB ID 9N83). A slightly variable kink angle allows accommodation of diverse DNA ends substrates. By contrast, the Lif1 helices remain straight, creating a rigid scaffold that restricts Dnl4_BRCTs_-Ku mobility (**Fig. 3a, 3b**) and generates an angle of ∼85° between the Nej1-Lif1 assembly and the DNA (**Fig. 3b**). The deviation of Dnl4_BRCT2_-Lif1 from Lig4_BRCT2_-XRCC4 is reflected in the altered orientation of the Nej1-Lif1 scaffold (**Fig. 3b**). These architectural differences translate into distinct DNA overall trajectories over the DSB. In the yeast complex, the DNA is bent by ∼135° (**Fig. 3a**), compared with ∼146° in the human assembly (**Fig. 3a**). In the centre of the ω-shaped SR synapsis complex, Nej1 contacts the N-terminal region of Lif1 through hydrogen bonds and hydrophobic contacts that involve conserved residues in yeasts (**Fig. 3c, Extended Data Fig. 7**). Lif1 stabilizes Dnl4 by interacting with its tandem BRCT domains through its extended α-helical arms across an extended ∼1000 Å² surface. This stabilization of Dnl4 involves both hydrophobic contacts and hydrogen bonds (**Fig. 3d**).

**Fig. 3.**
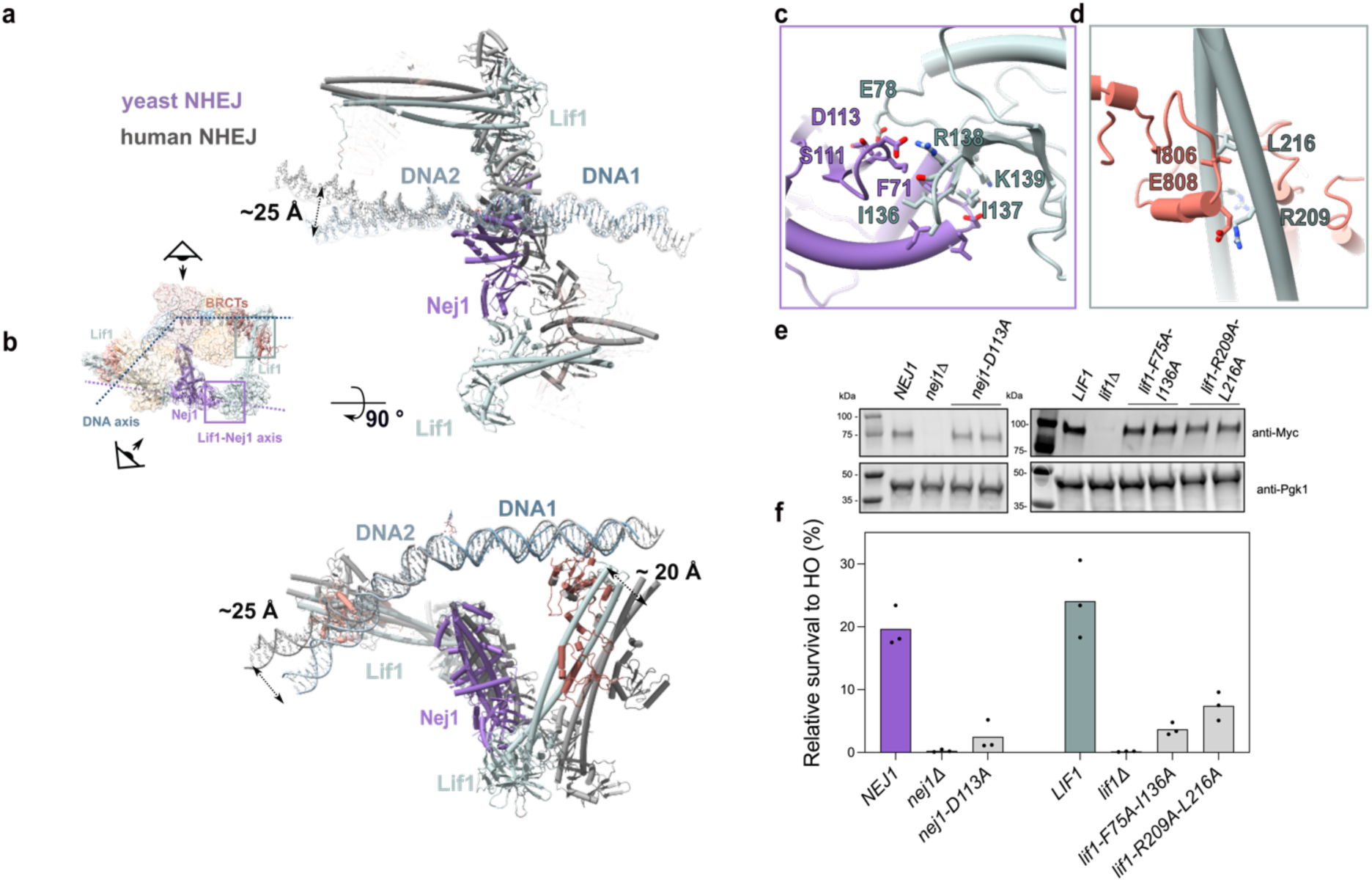
Differences in geometric constraints imposed on DNA conformation and orientation by Lif1/XRCC4 dimers in yeast and human through specific protein-protein interfaces. **a,** Comparison of the yeast end-joining complex and the human NHEJ complex (in grey PDB ID 9N83) after superposition of the catalytic modules of DNL4. Top view of the yeast NHEJ complex. The long coiled-coils of the Lif1-Nej1 assembly form a straight axis with an angle of 85° relative to the DNA axis, with no kink at the interface with the BRCT domains of Dnl4. In the human synaptic complex this angle is 60°. **b,** Side view. The long helix of Lif1 is shifted by ∼25 Å from the human XRCC4 corresponding position (distance measured between residues N233 and N200). **c,** The primary Lif1-Nej1 interface is mediated by key hydrogen-bonding involving D113^Nej1^ and R138^Lif1^, which led to the design of mutant D113A, and hydrophobic contacts between F71^Nej1^ and F75-I136^Lif1^ which were addressed by the double mutant F75A-I136A; other residues in the interface but not mutated in this study are also labelled. **d,** A second interface occurs between Lif1 and the BRCT domains of Dnl4, involving residues E868-I866^Dnl4^ and R209-L216^Lif1^, respectively. This interaction was addressed by the double mutant R209A-L216A. **e,** Western blot analysis of WT and mutant *Nej1* and *Lif1* proteins, all tagged with 13xMyc and detected using an anti-Myc antibody. **f,** Relative survival following HO expression, normalised to the uninduced condition.

To assess the contribution of the Nej1-Lif1 interface, we generated point mutations on either Nej1 (D113A) or Lif1 (F75A-I136A), targeting residues that stabilize the Nej1-Lif1 assembly (**Fig. 3c**). Following a pulse of HO endonuclease induction to produce a 4-bp microhomology overhang, both mutants displayed reduced end-joining efficiency relative to wild type, indicating that stabilization of this interface is essential for synapsis integrity (**Fig. 3e**). Since Lif1 can also be recruited via its interaction with the Dnl4 BRCT domains, we tested a *lif1*-*R209A-L216A* mutant allele, which disrupts this second interface (**Fig. 3d**) and observed a reduction in NHEJ efficiency. Importantly, those mutants do not grossly impact protein stability, consistent with the observed NHEJ defects being the consequences of impaired protein-protein interactions (**Fig. 3e**). We also generated a *lif1* quadruple mutant to disrupt both interfaces simultaneously (F75A-I136A-R209A-L216A), which resulted in a loss-of-function phenotype; however, in this case the combined mutations compromised protein stability, reducing Lif1 expression by approximately 80% (**Extended Data Fig. 6**). Together, these results support the conclusion that the Nej1-Lif1 assembly observed in the cryo-EM structure is essential for NHEJ *in vivo*.

### Differential engagement of two Dnl4 catalytic cores in two protective conformations

In yeast and human NHEJ, two Lig4 molecules are anchored to the synaptic complex through their BRCT domains bound to XRCC4/Lif1, forming a scaffold together with XLF/Nej1 ^25,26,56^. In the sequential model proposed for the human system, a single catalytic module (DBD-NTD-OBD) ligates one strand while the second module remains disengaged. After ligation of the first strand, the second module engages, the OBD closes, and the pre-adenylated Lig4 ligates the opposing strand ^17,24^.

In the first reconstituted budding yeast complex, the DNA substrate contained two different unligatable nicks: one 5’-phosphate/3’-dideoxy terminus (5’-P/3’-ddC) and an adjacent 5’-OH/3’-dideoxy terminus (5’-OH/3-’ddC) (**Fig. 4a**). Although both Dnl4 BRCT domains remained clearly visible in the cryo-EM density (**Extended Data Fig. 2**), only a minority (11 %) of particles displayed a fully ordered catalytic module (**Fig. 4a-c**). In this conformation, the module fully encloses the DNA, with clear density for AMP transferred to the 5′ phosphate at the nick (**Fig. 4c**). Additional low-resolution density was frequently observed near the second nick, suggesting the presence of a second DBD-NTD module that could not be modelled confidently. To clarify these additional conformational states, we studied another NHEJ complex with a DNA substrate containing two 5′-phosphate at the ends, which is the most likely biological substrate (**Fig. 4d**; **Extended data Fig. 3**). In order to trap the post ligation state of the first nick and observe the arrival of the second ligase, we used a modified 4-bp cohesive-end substrate containing one ligatable nick (5′-P/3′-OH) and one non-ligatable nick (5′-P/3′-ddC), an arrangement not previously examined by cryo-EM (**Fig. 4d**). Assembly was stabilized using GraFix (**Extended Data Fig. 3**). The structure was resolved to 3.3 Å overall, with local resolution of ∼3 Å for Ku and ∼4.5 Å for the Lif1-Nej1 complex. These components occupy positions comparable to those in the ligation-competent complex. The consensus map reveals two DBD-NTD modules of two Dnl4 molecules bound at the two nicks (**Extended Data Fig. 3**). Focused classification at the DNA DSB region resolved two predominant conformations comprising 68,870 and 62,686 particles, respectively (**Fig. 4e-g; Extended Data Fig. 3),** which we designate DNA-aligned protective conformations, with approximate reciprocal geometries.

**Fig. 4.**
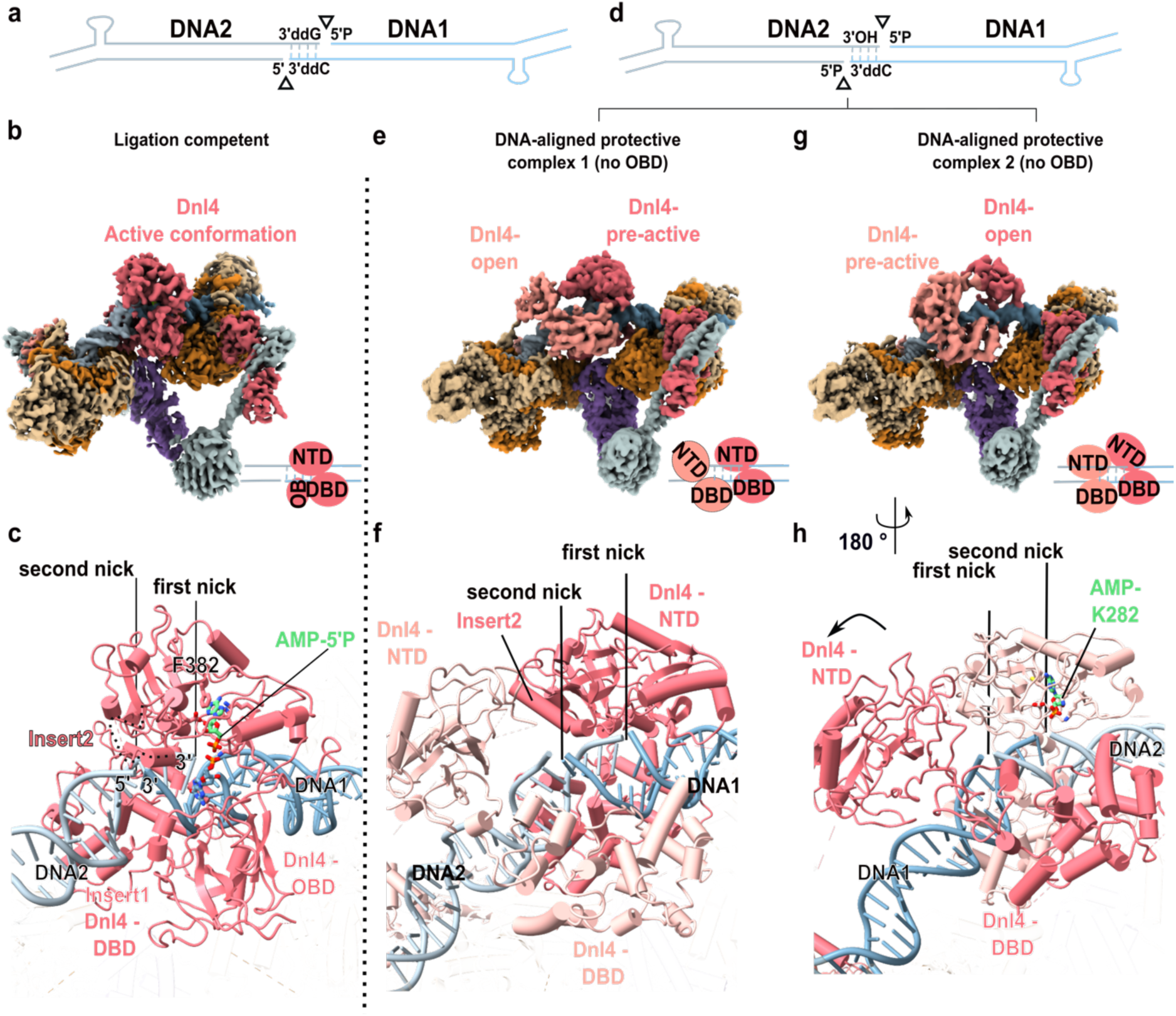
DNA substrate with two 5’-P at DNA ends reveals two additional conformations of the Dnl4 NTD-DBD domains indicative of an alternating ligation model. **a,** Schematic representation of the DNA substrate used to reconstitute the yeast NHEJ synaptic complex shown in **b** (same substrate as in Figs. 1-3), to be compared with **d**. **b,** Overall structure of a ligation-competent yeast NHEJ framework assembled on a cohesive-ended DNA substrate with a schematic representation of the ordered NTD-DBD-OBD module on a nick. **c,** With a DNA substrate containing a single 5′ phosphate (5′P), one of the conformations contains a Dnl4 molecule that adopts a closed conformation in which both the DBD and NTD engage the DNA end, while the OB domain assumes a closed, catalytically competent position, locked on the DNA, similar to that observed in the hNHEJ complex. The bottom panel shows the ligase catalytic site, with AMP molecule in ball and stick representation already migrated to the 5’-P. **d,** Schematic representation of an alternative DNA substrate carrying two 5′-phosphates adjacent to the DSB, yielding two other conformations shown in **e** and **g**. **e, g,** Cryo-EM structures of two distinct conformations (conformation 1 and conformation 2) of the yeast NHEJ complex assembled on 4-bp 3′ overhangs bearing two 5′ phosphates, one on each DNA end. The DBD domains remain at the same position on the DNA, but the NTD domains differ in orientation. **f, h,** In both conformations, two DBD-NTD modules of two different Dnl4 approach the DSB region. However, only one NTD is oriented in a geometry compatible with a pre- or post-ligation state, whereas the second NTD remains in an open conformation. Arrows indicate the pivoting motions of NTD domains around the DNA. Bottom panels show close-up views of the ligase catalytic site with the adenylated K282 sidechain in both conformations.

In the first conformation, the DBD-NTD module of one Dnl4 engages the ligatable nick (**Fig. 4f**). The OBD is not resolved (**Fig. 4f**) and the density does not allow modelling of an AMP moiety in the active site. The position of the DBD-NTD module and the distance between the 3’-OH and the 5’-P are compatible with a post-adenylation configuration, which we can only tentatively assign at this resolution. At the opposite nick, cryo-EM density is observed for the second Dnl4 DBD-NTD module. Although its DBD binds DNA similarly to the active state, the NTD is displaced by ∼25 Å relative to the pre-active position, suggesting only a remote engagement. Notably, Insert1 in both DBDs is disordered, indicating an inactive configuration.

In the second conformation, both DBD-NTD modules remain bound but adopt a reversed configuration relative to the first state (**Fig. 4g**). At the ligatable nick, the density is compatible with a direct link between the 3’-OH and the 5’-P and the NTD has pivoted away from the DNA (∼16 Å). Conversely, at the non-ligatable nick, the NTD of the second Dnl4 rotates by ∼25 Å to reach a pre-active configuration (**Fig. 4h**). Both DBD domains remain in place. However, OBD closure is not observed due to steric incompatibility with the opposing DBD domain. Concomitantly, the DNA duplex shifts relative to the complex, rotating by ∼39° and translating by ∼3.5 Å along its axis. This movement aligns the second nick with the catalytic module, and density corresponding to an AMP moiety covalently bound to K282 is visible (**Movie S1**). In addition, Insert1 is clearly resolved in this state, consistent with a pre-active configuration.

These last two conformations represent different intermediates along the ligation pathway. In the first population, the catalytic module bound at the ligatable nick adopts a configuration compatible with either a post-ligation state, while the second NTD remains displaced from the DNA. In the second population, the opposing catalytic module adopts a clear pre-active configuration; however, the first ligase remains positioned at the DSB, preventing OBD closure through steric hindrance. This structure therefore captures an intermediate in which the second catalytic module is in an interrogative mode (is everything in place to proceed further?) but cannot complete activation, due to the absence of a 3’-OH. The transition between the two states models the necessary rearrangements between the first and second ligation (**Movie S1**).

These observations support a sequential ligation mechanism in which the two Dnl4 catalytic modules alternately engage the DNA ends. Both DBD-NTD modules initially bind and align the DNA ends in a protective configuration. Ligation of the first strand requires displacement of the opposing DBD module to allow OBD closure and catalysis. Following this step, reopening of the OBD permits repositioning of the second catalytic module, enabling ligation of the remaining strand. Transient interactions between the NTD domains maintain the inactive ligase in proximity to the DNA ends, preserving end protection.

### Protecting and maintaining apart the DNA ends in a blunt-ended DNA substrate

To evaluate how Dnl4 handles other DNA end structures, a synapsis complex was assembled with a blunt-ended substrate lacking any microhomology region and carrying two 5′-phosphates and two 3′-ddC termini (**Fig. 5a; Extended Data Fig. 4**). Cryo-EM reconstruction of this complex, determined at 5.8 Å resolution, revealed two Dnl4 catalytic modules bound in an almost symmetric manner to the opposing 5′-phosphate termini (**Fig. 5b**). The overall architecture of Ku and Lif1-Nej1 matched that of the 4-nt overhang complex (**Fig. 5c**). In addition, the two DBD-NTD modules could be confidently docked into the map and this generates a major difference in the way the two DNA ends approach each other. Each 5′-phosphate is engaged in its NTD binding pocket, generating a symmetric intermediate in which the two DBD-NTD modules stabilize the ends but sterically hinder their DNA ends alignment by imposing a ∼30 Å distance between the 3’-OH and 5’-P ends (**Fig. 5d**). The second DBD-NTD bound at the DSB site is incompatible with OBD closure of the first DBD-NTD (**Fig. 5d**), because of a steric clash as observed before. Due to the distance between the two DNA ends, never observed before, and the peculiar way it protects the DNA ends, we call this state the non-aligned protective state, analogous to a medium-range synapsis complex. In the structure, residues 346-375 of Insert2 form a loop with a short 17-helix structure that is involved in the dimerization of Dnl4 catalytic modules (**Fig. 5d**). This sequence is present in the whole *Saccharomyces* clade (*Z. fermentati*, *K. lactis*, *C. glabrata*) and in *D. discoideum*. In contrast, the equivalent region in human Lig4 and in other species in the MSA is shorter and remains disordered (**Extended Data Fig. 1c–d**).

**Fig. 5.**
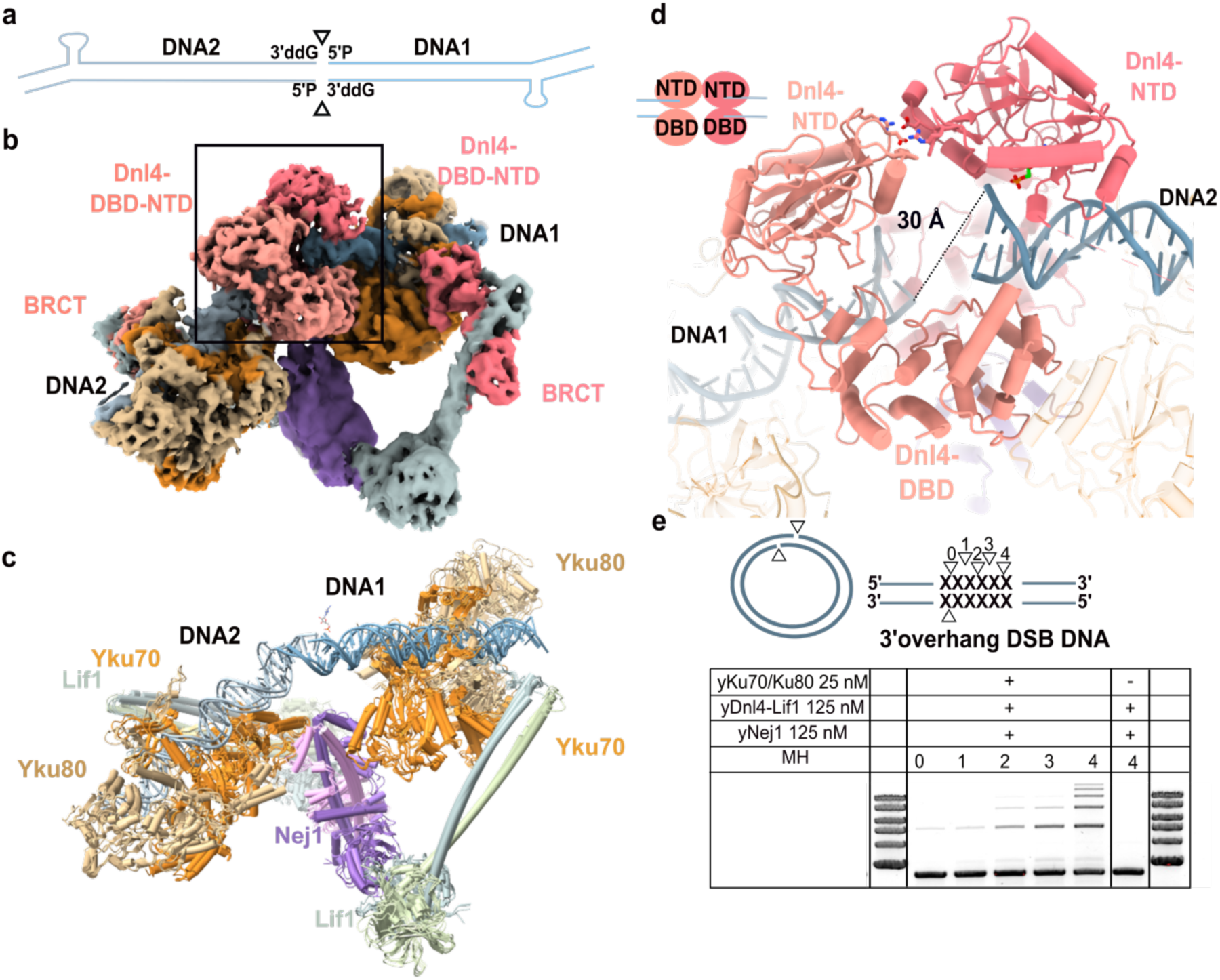
A blunt-ended DNA-DSB substrate reveals an unexpected protective and inactive synaptic complex allowing end processing. **a,** DNA substrate with a blunt ended DNA-DSB substrate carrying two 5’-P and 3’-ddG end modifications **b,** Cryo-EM structure of the corresponding yNHEJ complex. Ku70/80 and Nej1/Lif1 occupy positions similar to those observed in the cohesive-end complex. However, the two NTD domains of the ligases bind to the 5’-P ends and dimerize, protecting the DNA ends but separating them in an unusual protective but inactive synaptic complex. **c,** Structural superimposition of Ku-Lif1–Nej1 complexes bound to cohesive-ended and blunt-ended DNA substrates reveals identical positioning of these factors. For clarity, the DBD-NTD modules of Lig4 have been omitted. **d,** Zoomed-in view of both Dnl4 NTD engaging the DSB. The DNA termini are separated by ∼30 Å (dotted line) and are misaligned, preventing efficient ligation. Both NTD domains are engaged on a 5’-P and their dimerization mode prevents the DNA ends to be aligned in a productive way. **e,** Ligation assay. Various double-strand break (DSB) substrates derived from plasmid pUC19 were generated by restriction digestion to produce 4-, 3-, 2-, or 1-nt 3′ overhangs, as well as blunt ends. Dnl4 ligation was significantly more efficient with increasing microhomology length and not efficient with blunt ends, in accordance with the structural constraints seen in the cryo-EM structure.

A ligation assay was performed using substrates containing a microhomology region of varying sizes. The results indicated a clear correlation between microhomology length and ligation efficiency, with longer microhomology regions (2-4 bp 3’-overhang) supporting robust end-joining and not the short ones (1-bp 3’-overhang). For substrates with blunt-end termini, the ligation product was nearly undetectable, suggesting either very low ligation efficiency or significantly slower reaction kinetics (**Fig. 5e**). This protective and inactive synaptic complex observed in the cryo-EM structure of the blunt-ended synapsis is consistent with the *in vitro* ligation assays (**Fig. 5e**). In this configuration the blunt DNA ends are not aligned for ligation (**Fig. 5d**). This geometry suggests that the simultaneous engagement of both DBD-NTD protects the DNA ends but that the synapsis needs to undergo end processing and/or large rearrangements to become compatible with ligation. Assuming that the cryo-EM structure represents the most stable conformation of this complex in solution, productive repair of blunt ends will require the NTD of one ligase to transiently disengage or reposition to allow end alignment. This large rearrangement is likely to be slow, leaving ample time for a nuclease or a polymerase (Pol4) to intervene in order to create the possibility of short MH regions.

In summary, both *in vitro* and *in vivo* data in yeast point to an inefficient ligation reaction for blunt end DNA, and this is compatible with the structure shown in **Fig. 5** ^51^. This non-aligned protective complex that maintains the DNA ends widely separated, similar to the long-range complex observed in presence of DNAPKcs, but would necessitate substantial rearrangement to accommodate the action of a nuclease or a polymerase.

## Discussion

We determined cryo-EM structures of the yeast NHEJ machinery assembled on distinct DNA substrates, capturing ligation-competent, intermediate DNA-aligned and non-aligned protective synaptic complexes that are intermediates in the completion of double-strand ligation in a DNA-PKcs-independent system (**Fig. 6**). Although the overall architecture resembles mammalian short-range synapsis^17,24^ (state 2 in **Fig. 6**), the Lif1-Nej1 scaffold imposes a distinct DNA trajectory (**Fig. 3a**), with greater bending (135° in yeast versus 146° in human synapsis). Correspondingly, the angle between the DNA duplex and the Lif1- Nej1 scaffold increases from 60° to 85° (**Fig. 3b**), while the long α-helix of Lif1 remains straight at the Dnl4-BRCT interface. We propose that this rigid Nej1-Lif1-Dnl4^BRCT^ organization precisely positions the DNA termini for ligation and stabilizes two pre-positioned Dnl4 NTD-DBD catalytic modules, one at each nick. This arrangement gives rise to two aligned intermediates (states 1a and 1b; **Fig. 6**), in which one NTD-DBD module is engaged at the nick while the other remains remotely engaged, and vice versa (**Fig. 6**). The conformational transitions between intermediate states #1a and #1b of **Fig. 6** can be modelled by a movie (**Movie S1**), which illustrates the potential alternating activation of the two catalytic sites, allowing sequential nick ligation without disrupting the overall synaptic architecture. Concurrently, a coordinated rotation and translation of the DNA duplex, reorients each nick for optimal presentation to the active sites of Dnl4. In this process the Nej1-(Lif1-BRCT)_2_ complex moves very little. However, the Dnl4 catalytic core in its dimeric form is incompatible with productive closure of any of the two OBD. When one ligase catalytic core adopts an active, closed conformation, the second ligase catalytic core must leave the DNA break site. It appears therefore that in yeast ligation proceeds through alternating activation of two pre-positioned catalytic modules while the DNA ends remain continuously shielded within the synaptic complex.

**Fig. 6.**
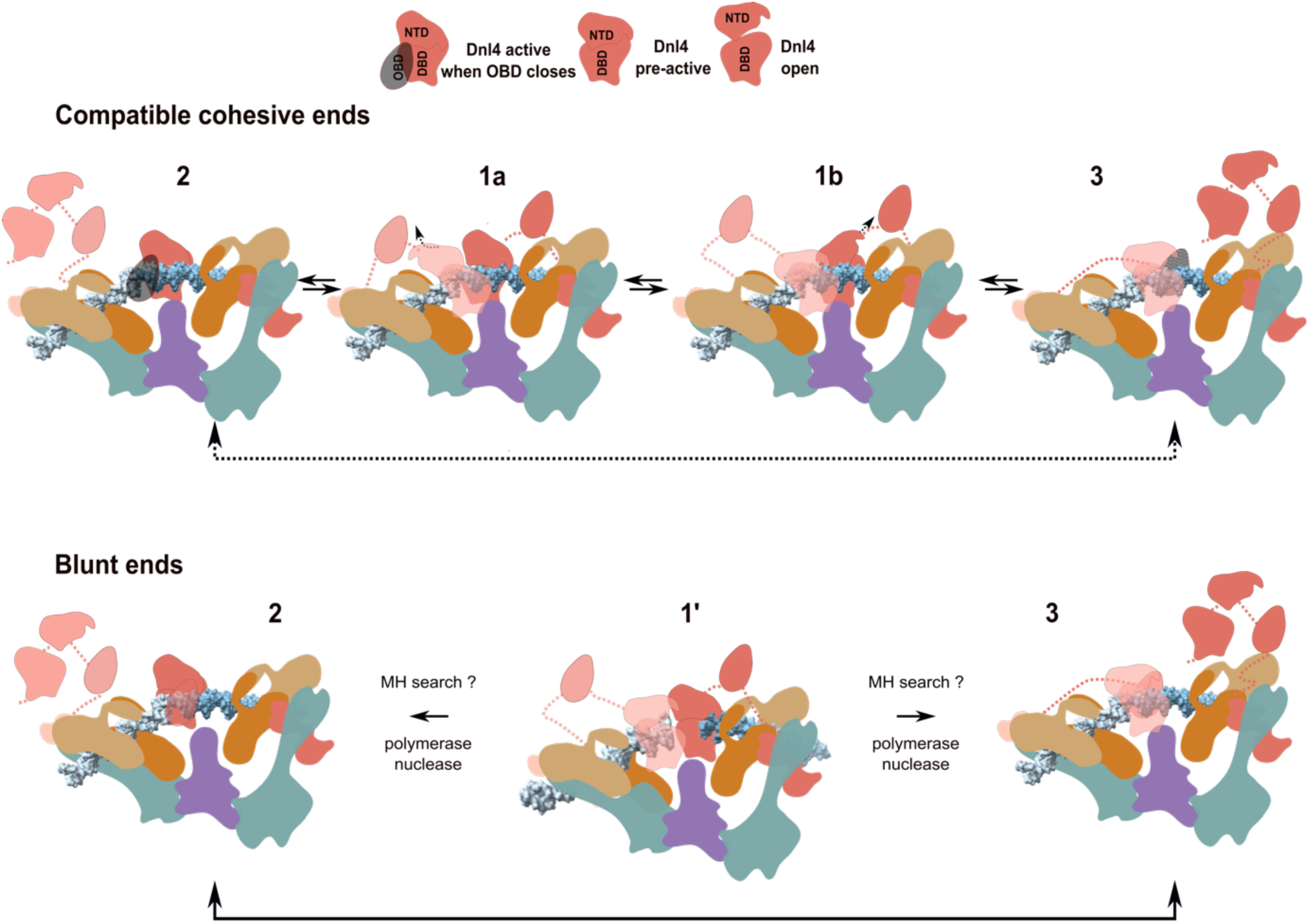
Proposed mechanism of ligation during end-joining in yeast. Top panel. In the case of cohesive-ends DNA breaks ready to be repaired with two 5’-P, the first ligation involves the **1a ⇔ 2** branch and the second ligation involves the 1b**⇔**3 branch. States #2 and #3 have one fully active ligase molecule, State #1a and 1b are protective states that interrogate alternatively the two DNA-ends to check if they are ligatable, with two DBD-NTD domains positioned on the DNA. In #1a the dark red NTD-DBD core is well positioned (on the right) and the pink NTD-DBD core (on the left) is on the verge to leave, in #1b it is the reverse. Bottom panel. For blunt-ended breaks, only one conformation #1’ was observed, in which both NTD-DBD modules of the two Dnl4 molecules engage the DSB simultaneously. The absence of any micro-homology region in the DNA and the interaction between the two NTD domains constrain the DNA ends to remain separated by ∼30 Å, a geometry incompatible with productive ligation. One NTD-DBD module must therefore vacate to permit closure of the OB domain and to allow the DNA ends to rotate and translate into an orientation in which the 3′-hydroxyl can attack the adenylated 5′-phosphate (states #2 or #3). Alternatively, nuclease and/or polymerase processing of the blunt-end DNA after the departure of at least one NTD domain could permit a transition to states #2 or #3, by finding a limited micro-homology region promoting access to the synaptic complex.

Our data provide a structural explanation for the slow reaction kinetics of the yeast core NHEJ machinery when processing blunt DNA ends ^51^. In this configuration, both NTD-DBD modules engage simultaneously, maintaining the DNA ends at a separation incompatible with productive chemistry. OBD closure therefore requires ejection of one NTD-DBD module and a coordinated DNA rearrangement (**Fig. 6**). Limited nucleotide addition or resection that generates minimal microhomology could relieve these constraints and promote entry into the DNA-aligned protective states observed with cohesive ends. The same scenario applies to NHEJ-dependent telomere-telomere fusions observed in yeast ^55,57^.

In the mammalian ligation-competent NHEJ synapsis complex, only one fully ordered and catalytically active Lig4 is resolved at the DNA break, whereas the second Lig4 is held only through BRCT-mediated interactions^17,24^. This may reflect either a more transient dual engagement of Lig4 in mammals or the presence of an alternative pathway for repair of the second nick^58,59^. Interestingly, the open conformation of human Lig4 captured in a gap-filling complex involving DNA polymerase µ resembles the open Dnl4 conformation observed in states #1a and #1b in yeast^24^. This suggests that such intermediates may be evolutionarily conserved, although the kinetics and regulatory context of catalytic domain exchange differ.

In addition, single-molecule analyses have shown that formation of the ligation-competent short-range complex in human cells is frequently preceded by reduction to a single Lig4 molecule^26^. Resolution of the second strand may therefore require rapid recruitment or re-engagement of a ligase, which is not necessarily Lig4. Indeed, DNA Ligase III (Lig3), typically associated with base excision repair (BER), has been proposed as an alternative source of ligase activity for second-nick sealing ^58–60^. Through its C-terminal domain BRCT Lig3 interacts with XRCC1, which in turn interacts with Pol β (a PolX with no BRCT domain), PARP, PNPK and Rev1^61^. However, *S. cerevisiae* and closely related saccharomycetes lack Lig3, XRCC1 and Pol β.

In DNA-PKcs-independent systems such as yeast, where in most cases the pol β-XRCC1-Lig3 is absent, the budding yeast system likely ensures completion of both ligation events within the same synaptic assembly. Without a rescue pathway, maintaining both DNA ends protected and pre-positioned becomes crucial. Conversely, in DNA-PKcs–dependent systems, such as vertebrates, delegation of second-nick ligation to Lig3-associated BER factors may reduce the necessity for a persistent dual-engagement protective state ^58,59^. The phylogeny distribution of DNA-PKcs is complex ^18^ but fungi do not have one in general, while most vertebrates have one (**Extended Data Fig. 1b**).

The protective complexes in the yeast NHEJ system described here are likely to be conserved in all <u>saccharomycetales</u> closely related to *S. cerevisiae*, such as *K. lactis* and *C. glabrata*. These species share the long Insert2 region implicated in NTD dimerization and second-nick protection (**Fig. 4f-h**, **Extended Data Fig. 1a, 1b**). Insert1 provides further evolutionary insight. In vertebrates (Type 2), a conserved threonine mediates a key hydrogen bond in XLF interaction. In *S. cerevisiae* (Type 1), this role is instead fulfilled by lysine or arginine shifted by two residues. More divergent fungi such as *S. pombe, S. japonicus, N. crassa*, and *A. clavatus* display vertebrate-like Insert1 features. Insert3, spatially complementary to Insert1, co-evolves accordingly. Phylogenetic analysis of Lig4 sequences recapitulates these structural distinctions (**Extended Data Fig. 1b, 1c**).

Parallel evolutionary trends are observed in PolX family members ^62,63^. Some fungi possess Pol µ-like enzymes (*S. pombe, S. japonicus*), others Pol λ-like enzymes (*G. zeae*), and some harbour both (*N. crassa, A. clavatus*). Notably, *G. zeae* and related species encode a putative Lig3, consistent with their vertebrate-like Insert1 features. These observations support the idea that BER components, particularly Pol β, emerged after divergence of the saccharomycete lineage (**Extended Data Fig. 1b**).

In conclusion, structural studies of the NHEJ system in yeast illuminate how a primitive system could work in the absence of DNA-PKcs. It takes advantage of stable protective states that lead to a complete ligation on both strands in most situations, with intermediates that involve the dimerization of the DBD-NTD catalytic core of the two ligases. It is possible that this dimer also exist in human very transiently, in which case it would constitute a new target for drug design since trapping it would block the reaction. Finally, a possible explanation of the success of the human NHEJ system that contains more added factors than budding yeast is that it can delegate the second ligation to an auxiliary pathway in DNA repair (BER).

## Acknowledgements

We thank Ludovic Sauguet and Marcin Suskiewicz for helpful discussions and suggestions. We thank Ludovic Deriano and Benoit Arcangioli for careful reading of the manuscript. We acknowledge SOLEIL and the French EQUIPEX+ France Cryo-EM (ANR-21-ESRE-0046) for provision of beam time at their cryo-EM facilities and we thank Heddy Soufari and Eric Larquet for assistance in using the POLARIS microscope under proposal number 99250139. SM was initially supported by ANR grant “DNA Break-Dance” 20-CE11-0026-03 (MD) and then by a grant from Fondation pour la Recherche Medicale, Equipe Labellisée 2024 03018410 to MD. This work benefited from the facilities and expertise of the Prot-EX platform at I2BC, supported by the French Infrastructure for Integrated Structural Biology (FRISBI, ANR-10-INBS-05). We thank Soizick Lucas-Staat (IP) for the MALDI-TOF experiments. MM and JBC were supported by ANR-20-CE11-0026 and ANR-23-CE11-0039-01.

## Contributions

Conception and design: SMy, SMd, MM, JBC and MD. Acquisition of data: SMy, CT, VR, PL, VM, PFV, SC, SMi, AP and MM. Analysis of data: SMy, PL, SC, SMi, SMd, MM, JBC and MD. Drafting the article: SMy and MD. All authors edited and approved the manuscript.

## Deposition

The ligation-competent complex has been deposited in the Electron Microscopy Data Bank (EMDB) under accession number EMD-XXXX and in the Protein Data Bank (PDB) under accession number YYYY.

The DNA-aligned protective states 1a and 1b complexes have been deposited under accession numbers EMDB-XXXX and PDB-YYYY and EMDB-XXXX and PDB-YYYY, respectively. The non-aligned protective state complex has been deposited under accession numbers EMDB-XXXX and PDB-YYYY.

## Material and Methods

### Proteins expression in insect cells

Recombinant yeast NHEJ factors were produced using the baculovirus expression system. Codon-optimized genes encoding Lif1-Dnl4 and Yku70/Yku80 were cloned into the pKL vector while Nej1 was cloned into the pFL vector (synthesized and cloned by Genscript). Recombinant bacmids were generated using the MultiBac system. Baculoviruses were amplified once and used to infect Sf21 cells cultured at 27 °C in Sf-900^TM^ II SFM medium (Gibco) at a density of 2-2.5 × 10⁶ cells ml⁻¹. Cells were harvested 48-60 h post-infection, washed with PBS 1X, flash-frozen in liquid nitrogen, and stored at −80 °C.

### Protein purification

1. Cell pellets of <u>Lif1-Dnl4</u> were thawed on ice and resuspended in lysis buffer (20 mM Tris pH 8.0, 150 mM NaCl, 150 mM KCl, 20 mM imidazole, 5% glycerol, 0.5mM TCEP) supplemented with proteases inhibitors cocktail tablets (*Roche*). Cells were lysed by sonication and clarified by centrifugation at 20,000 rpm for 20 min at 4 °C and mixed with 4 µL benzonase (*Sigma, 25kU*). Supernatants were loaded onto Ni-NTA resin (*Macherey-Nagel*), washed with the lysis buffer supplemented with 850mM NaCl, and eluted with lysis buffer containing 350 mM imidazole. Eluted fractions were diluted, applied to a Resource Q anion-exchange column (*Cytiva*) equilibrated in buffer A (50 mM Tris pH 8.5, 100 mM NaCl, 5 mM EDTA, 0.5 mM TCEP). Proteins were eluted with a salt gradient from 0 % to 40% buffer B (20 mM Tris pH 8.0, 150 mM KCl, 850 mM NaCl, 0.5 mM TCEP). Peak fractions were concentrated to 3 mg/ml, dialysed in 50 mM Hepes pH 7.9, 100 mM KCl, 5% glycerol, 0.5 mM TCEP) flash-frozen and stored at -80°C (**Extended Data Fig. 5**). Protein yield is 8.91 mg per liter of insect cells culture.
2. Cell pellets of <u>Nej1</u> were thawed on ice and resuspended in lysis buffer (20 mM Tris, pH 8.0, 150 mM NaCl, 150mM KCl, 20 mM imidazole, 5% glycerol) supplemented with proteases inhibitors cocktail tablets (*Roche*). Cells were lysed by sonication, clarified by centrifugation at 20,000 rpm for 20 min at 4 °C and mixed with 4 µL benzonase (*Sigma, 25kU*). Supernatants were loaded onto Ni-NTA resin (*Macherey-Nagel*) resin, washed with lysis buffer supplemented by 850 mM NaCl and eluted with lysis buffer containing 300 mM imidazole. Eluted fractions were diluted, applied to a Resource Q Sepharose anion-exchange column (*Cytiva*) equilibrated in buffer A (50 mM Tris pH 8.5, 50 mM NaCl, 50 mM KCl, 0.5 mM TCEP). Proteins were eluted with a 0 % to 40% buffer B (20 mM Tris pH 8.0, 150 mM KCl, 850 mM NaCl, 0.5 mM TCEP) salt gradient. Peak fractions from the Resource Q were concentrated to 3 mg/ml and dialysed in 50 mM Hepes pH 7.9, 150 mM NaCl, 5% glycerol, 0.5 mM TCEP), flash-frozen and stored at -80°C. (**Extended Data Fig. 5**). Protein yield is 9.97 mg per liter of insect cells culture.
3. Cell pellets of <u>Ku</u> were thawed on ice and resuspended in lysis buffer (20 mM Tris pH 8.0, 150 mM NaCl, 50 mM KCl, 0.1% Triton, 5% glycerol, 0.5 mM TCEP) supplemented with protease inhibitors cocktails proteases tablets (*Roche*). Cells were lysed by sonication and clarified by centrifugation at 20,000 rpm for 20 min at 4 °C, then incubated with 4 µL benzonase (*Sigma, 25kU*). Supernatants were loaded onto Ni-NTA resin (*Macherey-Nagel*), washed, and eluted with 300 mM imidazole linear gradient. Eluted fractions were diluted, applied to a Resource Q anion-exchange column (*Cytiva*) equilibrated in buffer A (20 mM Hepes pH 8.0 and 50 mM NaCl, 50 mM KCl, 5 mM EDTA, 0.5 mM TCEP). Proteins were eluted with a 0 % to 100 % buffer B (20 mM Tris pH 8.0, 850 mM NaCl, 50 mM KCl, 0.5 mM TCEP) salt gradient. Peak fractions were concentrated to 2 mg/mL. Proteins were dialysed in 50 mM Hepes pH 7.9, 150 mM KCl, 0.5 mM TCEP, 5% glycerol, flash frozen and stored at -80°C. (**Extended Data Fig. 5**). Protein yield is 1.58 mg per liter of insect cells culture.

### DNA substrates design and assembly

Cohesive-end and blunt-end DNA substrates were prepared by annealing synthetic oligonucleotides followed by PAGE or ion-exchange purification (*Eurogentec*). All the sequences of oligonucleotides used for assemblies are summarized into the **Supplementary Table 2**. Duplexes were annealed into 10 mM HEPES pH 7.9, 50 mM KCl and 10 mM MgCl₂.

### GraFix assembly of end-joining complexes

End-joining assemblies were prepared by the GraFix method^64^ to stabilize the fragile NHEJ complexes. Protein-DNA mixtures were assembled on ice in 50 mM HEPES pH 7.9, 100 mM KCl, 5 mM β-mercaptoethanol and 5 mM Mg(OAc)₂. Glycerol gradients were prepared with a top buffer containing 10% glycerol and a bottom buffer containing 40% glycerol and 0.01% glutaraldehyde, both made in the same base buffer. A duplicate gradient lacking glutaraldehyde was run in parallel. Samples were centrifuged at 100,000 × g for 12 h at 4 °C. Fractions from the non-cross-linked gradient were analysed by SDS-PAGE to identify those containing the intact assemblies. The corresponding fractions from the cross-linked gradient were collected, quenched with 1 µl Tris pH 7.5, dialyzed into 50 mM HEPES pH 7.9, 100 mM KCl, 5 mM β-mercaptoethanol and 5 mM Mg(OAc)₂, and concentrated to 0.3-1 mg ml⁻¹.

### Cryo-EM grid preparation

Complexes (3-3.5 µL) were applied at the top and the bottom to UltrAuFoil R1.2/1.3 holey gold grids glow-discharged for 35 s. Grids were blotted for 2 s at blot force 0, 4 °C and 100% humidity in a Vitrobot Mark IV and vitrified in liquid ethane. Grids were then stored in liquid nitrogen until collection. Grids were visualized on a 200 kV Glacios microscope to check the repartition of specimens on grids and a small data collection was done to evaluate samples quality.

### Cryo-EM data collection

All datasets were acquired on the POLARIS beamline at synchrotron SOLEIL (Saint Aubain, France) on a Titan Krios G4 microscope operating at 300 kV equipped with a Falcon 4i detector and a Selectris X energy filter set at 10 eV. Movies were recorded at a nominal magnification of ×130,000 in EER format, yielding a pixel size of 0.96 Å/pixel. A total dose of ∼40 or 50 e⁻ Å⁻² was delivered over 40-50 frames using automated acquisition schemes with a defocus range of -0.4 to -2.4 µm.

### Cryo-EM image processing

The detailed protocol is described below for the first complex carrying only one 5’-P. Image processing was performed using cryoSPARC^65^. Movies were corrected for beam-induced motion and gain variations using Patch Motion Correction, and contrast-transfer parameters were estimated with Patch CTF. Particle coordinates were identified using a combination of blob picking, template-based picking, and Topaz neural-network picking. After extraction, particles were iteratively cleaned through multiple rounds of 2D classification prior to ab initio reconstruction and heterogeneous refinement.

An initial consensus reconstruction revealed density corresponding to two Ku heterodimers, a Nej1 homodimer, and a Lif1 homodimer. However, the density near the DNA break, where the catalytic modules of Dnl4 are expected, was initially weak. Although the resolution was insufficient for confident model-building, low-resolution features suggested the presence of two distinct catalytic modules of Dnl4. To address conformational and compositional heterogeneity, focused 3D classification was performed using a soft mask surrounding the catalytic domains of Dnl4, enabling separation of discrete substrate-bound states and catalytic configurations. The best-resolved classes were subsequently refined using non-uniform refinement. Local refinements were performed independently for the Ku70-Ku80-DNA module, the Nej1-Lif1 filament (refined with C2 symmetry), and for DBD-NTD-OB set of Dnl4. Resulting local maps were further improved using Phenix Resolve, and local resolution estimates were obtained with Phenix tools^66^. Composite maps were generated with phenix.combine_maps to obtain the final representation of the end-joining assembly (**Extended Data Fig. 2**). Resolution estimation was performed by combining focused half-maps using the vop maximum tool in ChimeraX^67^ and phenix local resolution analysis tool ^68^.

For the remaining complexes (4-bp 3’-overhang carrying two 5’-P and blunt-end DNA substrates), a similar cryo-EM data-processing pipeline was applied, and the results are summarized in **Extended Data Figs. 3 and 4**.

### Model building and refinement

Initial models of Nej1 and Lif1 were obtained from AlphaFold3 ^69^ predictions and rigid-body docked into the corresponding cryo-EM densities using UCSF ChimeraX ^67^. Yeast Ku70-Ku80 was placed using the existing crystallographic model (PDB ID 5Y58). The BRCT domains of Dnl4 were placed using the available crystal structure (PDB ID 1Z56), and the catalytic region of Dnl4 was initially modelled using AlphaFold3. The DBD, NTD, and OB-fold domains were docked separately to optimize their positions and rigid body refinement with Namdinator server ^70^. Model adjustments were carried out manually in Coot ^71^, guided by local map features and secondary-structure predictions. Real-space refinement was performed in Phenix ^66^ using secondary-structure restraints, non-crystallographic symmetry restraints where applicable, and optimized weightings. Model geometry was evaluated using MolProbity ^72^ and EMRinger ^73^ to ensure stereochemical quality and consistency with the cryo-EM density. Data collection and model refinement statistics are reported in **Extended Data Table 1**.

### Movie generation

Structural transition movies were generated using UCSF ChimeraX. Atomic models corresponding to the different conformational states were superimposed and interpolated using the morph command, and animations were recorded using the built-in movie tools.

### In vitro ligation assay

To assess ligation activity, pUC19 plasmid DNA was linearized with a restriction enzyme to generate defined ends, gel purified and stored at 50 ng/µL in 10 mM Tris-HCl pH 8, 1 mM EDTA. Reactions (10 mL) contained 5 ng/µL of linear DNA, the indicated concentrations of Ku70/80, Nej1 and Dnl4/Lif1, 6 mM Tris-HCl pH 7.5, 0.3 mM EDTA, 3 % glycerol, 1.3 mM DTT, 2 mM MgCl2 and 45 mM KCl. Reactions were incubated at 30°C for 60 minutes and stopped by addition of 2 µL of 6X NEB purple loading buffer containing SDS, 2 µL of Pronase at 20 mg/ml and heated at 55°C for 20 min. Reaction mixes were fractionated by electrophoresis on 0.8% agarose gels in 1X TBE buffer at 80 volts for 90 minutes (15 cm wide, 10 cm long, 100 ml of gel). NEB 1 kb ladder was used as size marker. Gels were stained with GelRed (Biotium) in 1X TBE buffer for 20 minutes and rinsed with milliQ water for 2 hours. The fluorescent signal was captured with a BioRad ChemiDoc system.

### Strains and Plasmids

Strains and plasmids used in this study are listed in **Supplementary Table 3, 4.** All experiments were performed in haploid strains in which the *HO* endonuclease is expressed from a galactose-inducible promoter. To prevent repair by homologous recombination, *HML* and *HMR* were deleted. Plasmids expressing wild-type or mutant versions of myc-tagged Lif1, Nej1 and Yku80 were generated by transforming yeast strains lacking the endogenous gene with a linearized pRS315 centromeric plasmid (*LEU2)*, together with PCR fragments corresponding to the desired allele. Plasmids recovered from transformants were verified by sequencing, and expression of the mutant proteins was confirmed by Western blotting using extracts from exponentially growing cells.

### Yeast Survival Assays

Cells from strains Lev348 (WT), Lev379 (*lif1Δ*), Lev406 (*nej1Δ*), and Lev370 (*yku80Δ*), constructed and designed as described previously ^41^, were transformed with plasmids expressing either the wild-type or mutant allele, or with empty pRS315 as a control. Transformants were grown exponentially in raffinose-containing medium lacking leucine and synchronized in G1 with α-factor (10^−7^ M) for 3 h. A fraction of each culture was induced for *HO* expression by addition of 2% galactose for 45 min; parallel aliquots were incubated with 2% glucose as uninduced control. Cells were washed and resuspended in glucose- and α-factor-containing medium for 1 h to repress *HO* and allow accurate repair of the DNA break, then plated at appropriate dilutions on glucose medium lacking leucine. After 3 days at 30 °C, colonies were counted manually. Survival rates were calculated as the ratio of colony numbers on induced versus uninduced control plates. For assays involving continuous *HO* expression, exponentially growing cells were plated at serial dilutions onto galactose-containing medium lacking leucine.

