## Extended Data for "Structural basis for continuous DNA-end protection during ligation of double-strand breaks in yeast Non-Homologous End-Joining"

**a**

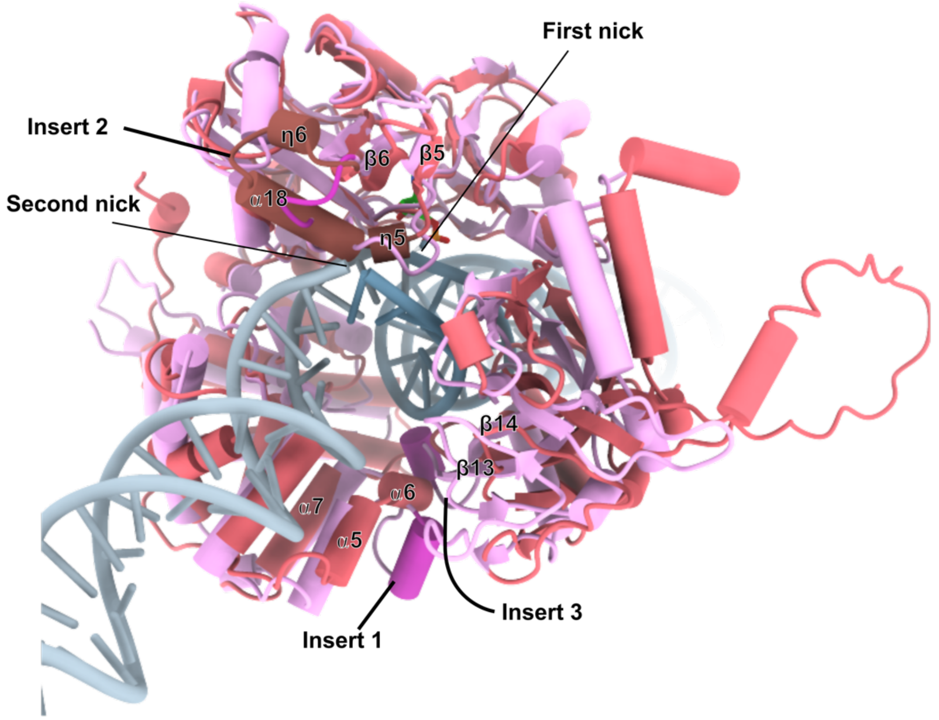


**b**

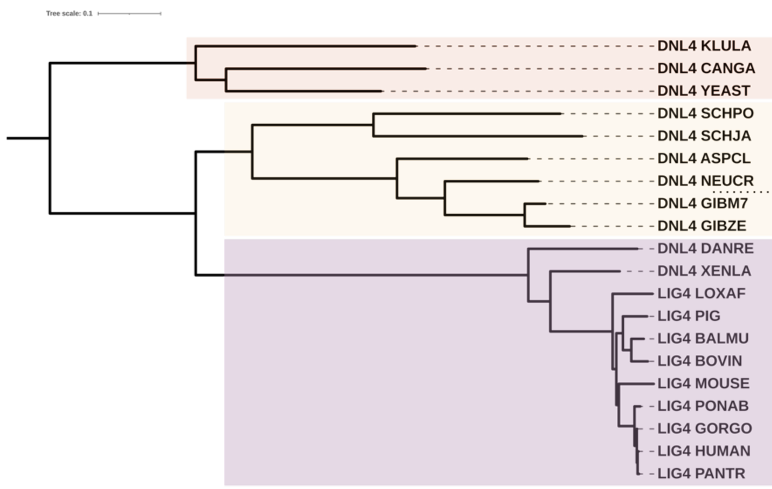


| | Insert 1 | Insert 2 | Insert 3 | DNA PKcs | LIG3 |
|---|---|---|---|---|---|
| DNL4 KLULA, DNL4 CANGA, DNL4 YEAST | Type 2 | Long | Short | No | No |
| DNL4 SCHPO, DNL4 SCHJA, DNL4 ASPCL, DNL4 NEUCR | Type 2 | Long | Short | No | No ? |
| DNL4 GIBM7, DNL4 GIBZE | Type 1 like | Long | Short | No | Yes ? |
| DNL4 DANRE, DNL4 XENLA, LIG4 LOXAF, LIG4 PIG, LIG4 BALMU, LIG4 BOVIN, LIG4 MOUSE, LIG4 PONAB, LIG4 GORGO, LIG4 HUMAN, LIG4 PANTR | Type 1 | Short | Long | Yes | Yes |

c

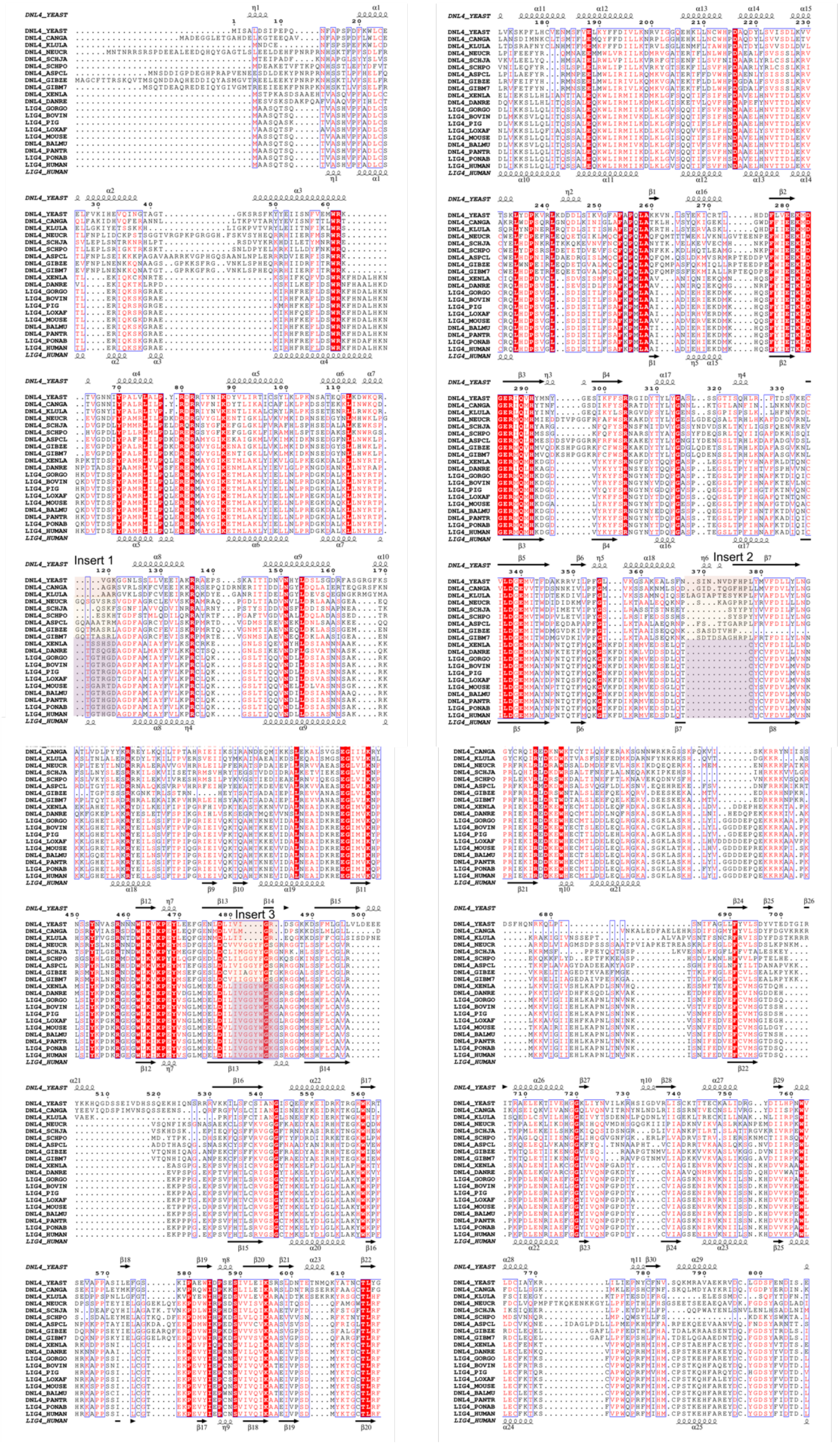

**Extended Data Fig. 1| Evolution of Dnl4**

**a,** Structural superposition of AlphaFold [1] model of *Saccharomyces cerevisiae* Dnl4 catalytic module (DBD-NTD-OBD domains) in brownish red with the PDB structure 9N83 of human LIG4 in pink (DBD-NTD-OBD domains) bound to a DNA DSB substrate. yDnl4 adopts an overall fold similar to that of hLIG4 (RMSD = 0.988 Å over 278 aligned residue pairs), but differs in the following regions: Insert1 (between α5 and α7), Insert2 (between β5 and β6), and Insert3 (between β13 and β14). These regions are highlighted in pink in the human LIG4 structure and in maroon red in yeast. Insert1 and Insert3 are close in space and their length is anticorrelated across species. Insert 2 loop (residues 362-375) is conserved within *Saccharomycetes* clade and close in space to the second nick in the DNA.

**b,** Phylogenetic tree based on sequence similarity generated using the iTOL server [2]. Insertions in each species were analyzed and correlated with the presence or absence of DNA-PKcs and DNA ligase 3 (LIG3) across evolutionary lineages.

**c,** Multiple sequence alignment (MSA) using ESPRIPT [3] of DNL4/LIG4 spanning the C-terminal domain (CTD) through the OBD. Sequence differences in Insert 1, Insert 2, and Insert 3 are highlighted in light maroon for the *Saccharomycetes clade*, orange for the *Schizosaccharomyces clade*, and pink for vertebrates, e.g. *Xenopus laevis, Danio rerio, and mammals*.

**Species abbreviations:** YEAST (*Saccharomyces cerevisiae*), CANGA (*Candida glabrata*), KLULA (*Kluyveromyces lactis*), NEUCR (*Neurospora crassa*), SCHJA (*Schizosaccharomyces japonicus*), SCHPO (*Schizosaccharomyces pombe*), ASPGL (*Aspergillus glaucus*), GIBZE (*Gibberella zeae*), GIBM7 (*Gibberella moniliformis*), XENLA (*Xenopus laevis*), DANRE (*Danio rerio*), GORGO (*Gorilla gorilla*), BOVIN (*Bos taurus*), PIG (*Sus scrofa*), LOXAF (*Loxodonta africana*), MOUSE (*Mus musculus*), BALMU (*Balaenoptera musculus*), PANTR (*Pan troglodytes*), PONAB (*Pongo abelii*), HUMAN (*Homo sapiens*).

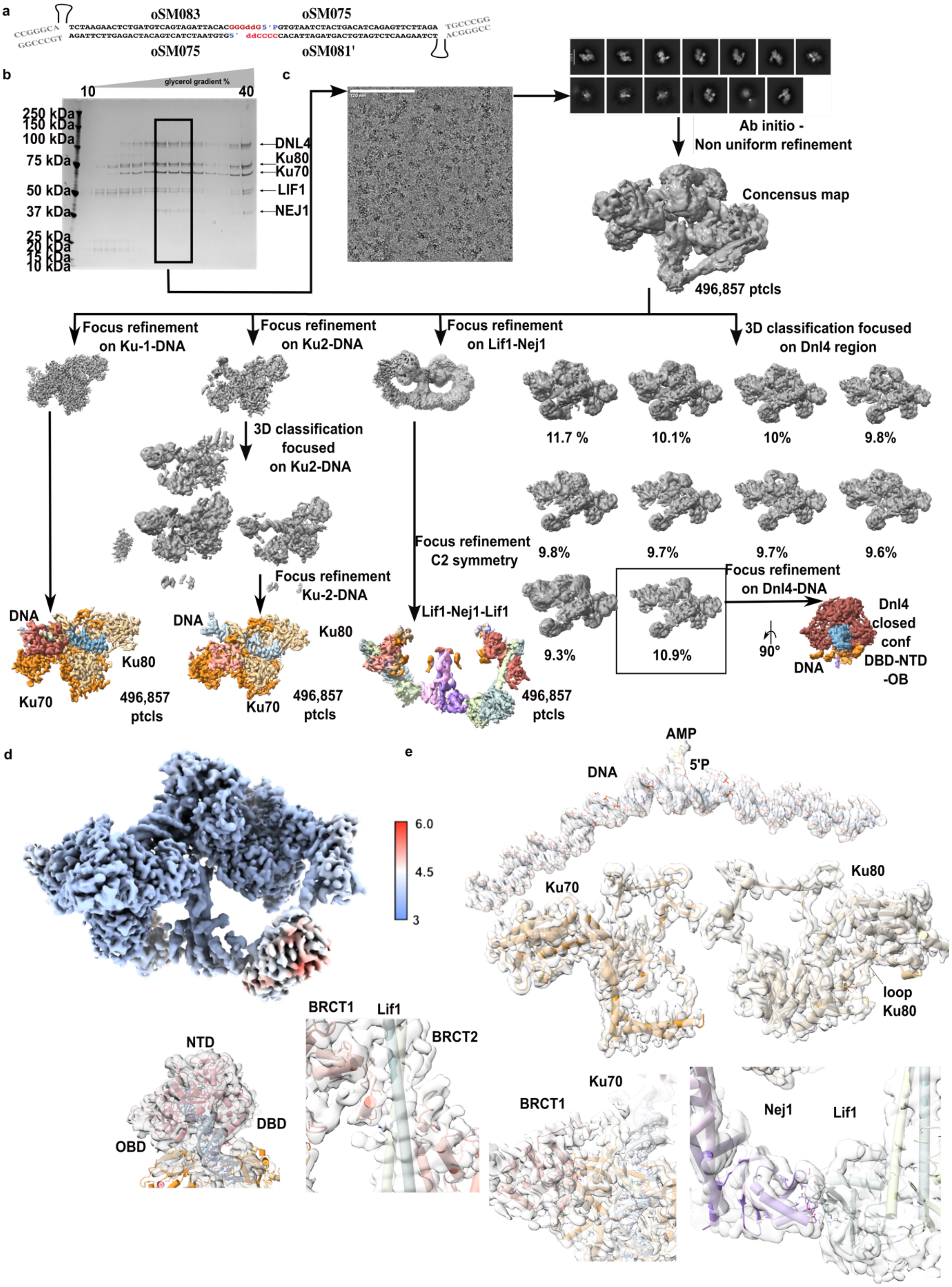


**Extended Data Fig. 2| Data processing of the ligation-competent end-joining complex**
**a,** DNA substrate. **b,** SDS–PAGE analysis of the GraFix-stabilized end-joining complex in the presence of the DNA substrate shown in **a**. **c,** Representative cryo-EM micrograph from 17,117 collected images and image-processing workflow. **d,** Composite cryo-EM map colored according to local resolution. **e,** Cryo-EM density of each region of interest.

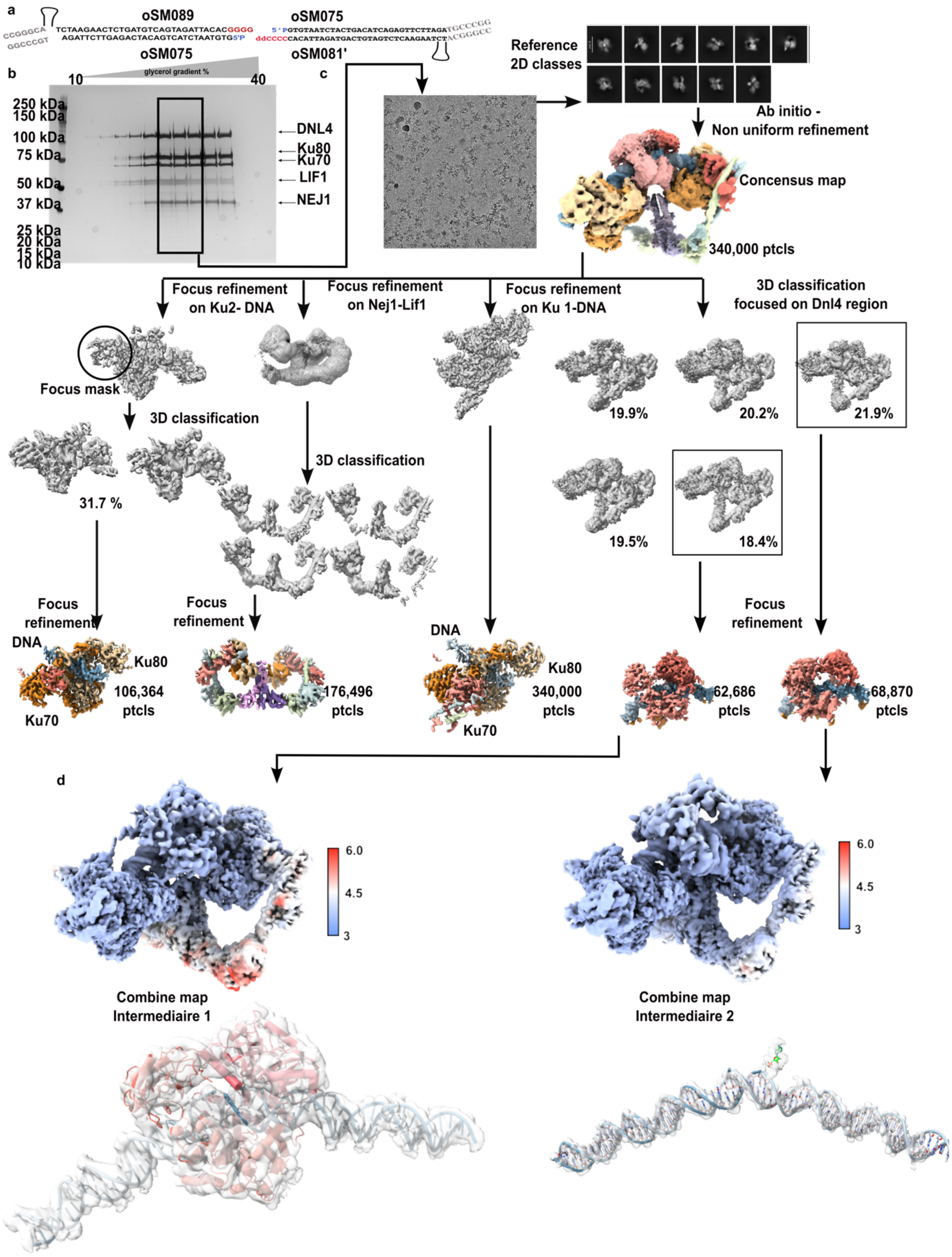


**Extended Data Fig. 3| Data processing of DNA-aligned protective states end-joining complex**
**a,** DNA substrate. **b,** SDS–PAGE analysis of the GraFix-stabilized end-joining complex in the presence of the DNA substrate shown in **a**. **c,** Representative cryo-EM micrograph from 12,350 collected images and image-processing workflow. **d,** Composite cryo-EM maps colored according to local resolution (top) and focused on one nick or on the DNA alone (bottom).

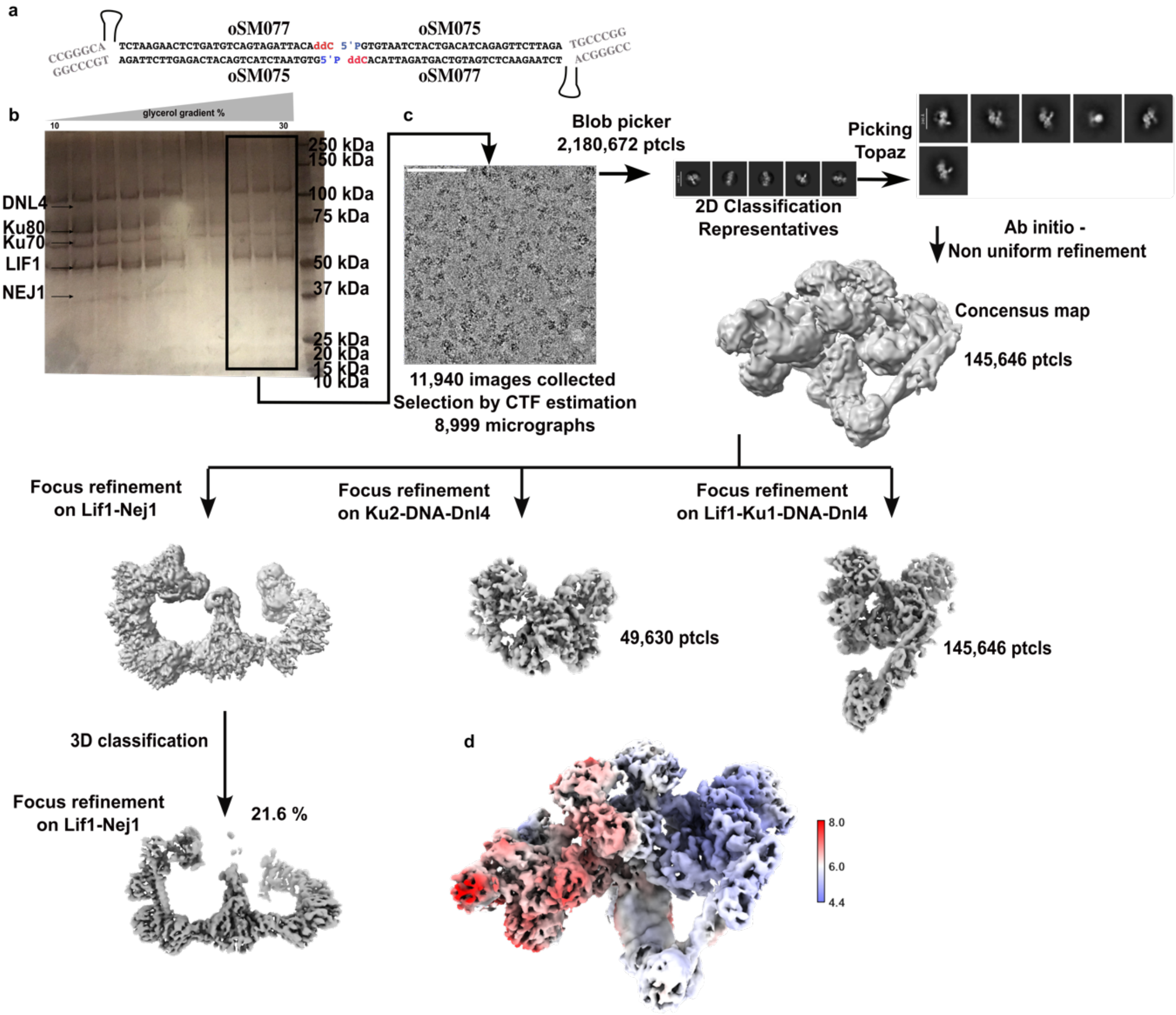


### Extended Data Fig. 4| Data processing of DNA non-aligned protective states end-joining complex

**a,** DNA substrate. **b,** SDS–PAGE analysis of the GraFix-stabilized end-joining complex in the presence of the DNA substrate shown in **a**. **c,** Representative cryo-EM micrograph from 11,940 collected images and image-processing workflow. **d,** Composite cryo-EM map colored according to local resolution.

a

b

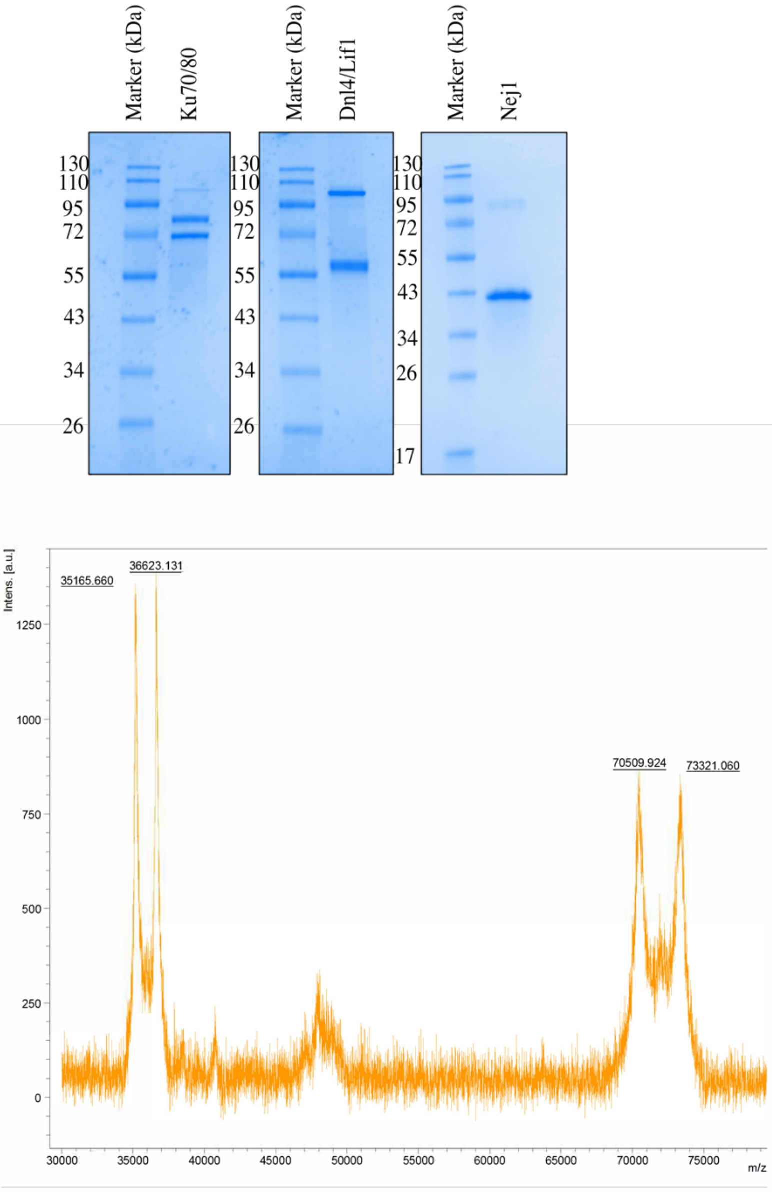

Marker (kDa)
Ku70/80
Marker (kDa)
Dnl4/Lif1
Marker (kDa)
Nej1
130
110
95
72
55
43
34
26
17
Intens. [a.u.]
35165.660
36623.131
70509.924
73321.060
1250
1000
750
500
250
0
30000
35000
40000
45000
50000
55000
60000
65000
70000
75000
m/z

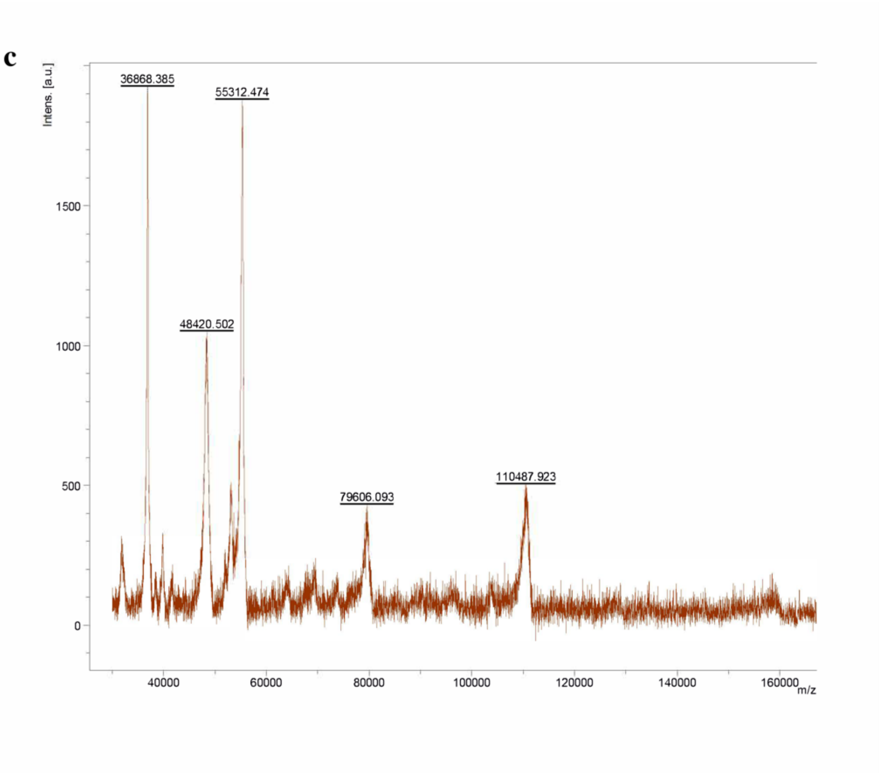


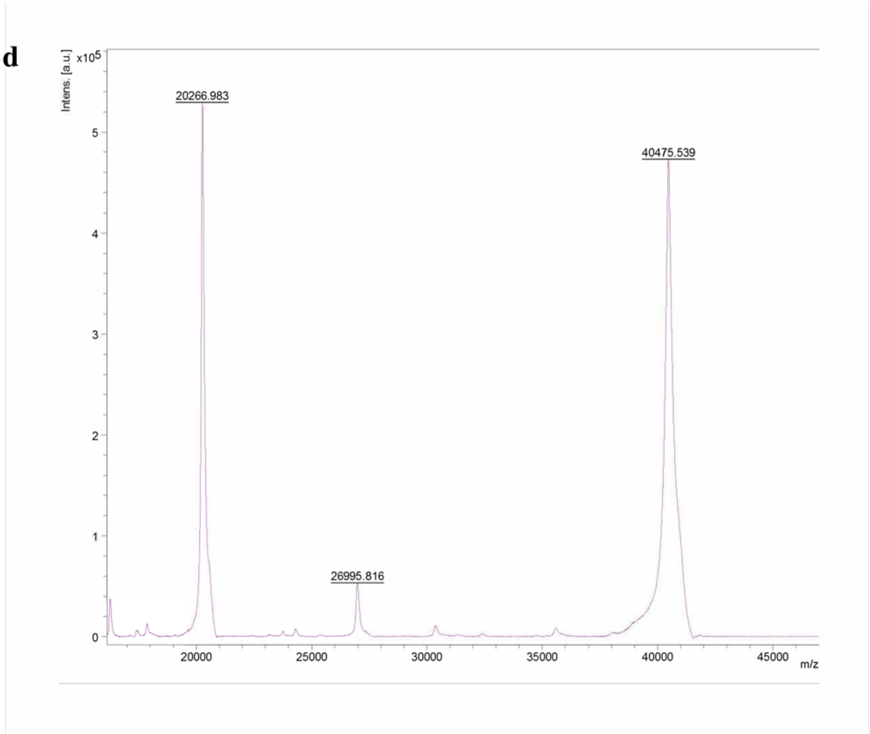


**Extended Data Fig. 5| Proteins quality assessment**
**a,** SDS PAGE of Ku70/80, Dnl4/Lif1 and Nej1.
**b,** Maldi-TOF mass spectrum of Ku70/80.
**c,** Maldi-TOF mass spectrum of Dnl4/Lif1.
**d,** Maldi-TOF mass spectrum of Nej1.

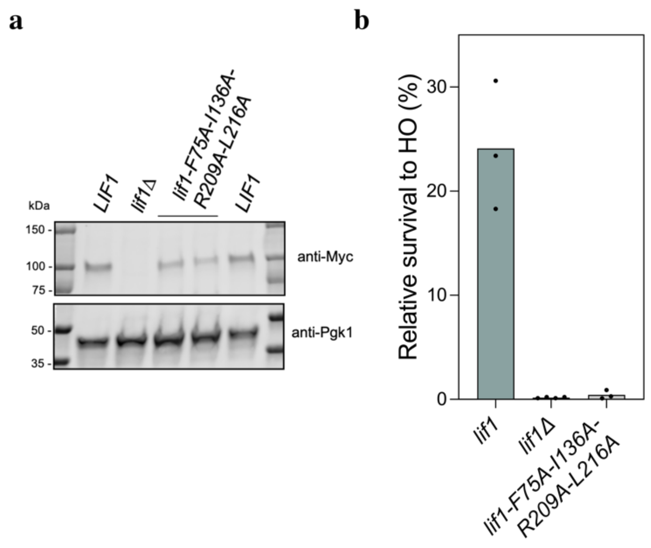


**Extended Data Fig.6| End joining assay after a transient HO pulse of mutant allele *lif1-F75A-I136A-R209A-L216A*.**

**a,** Western blot of LIF1, *lif1Δ, lif1-F75A-I136A-R209A-L216A* alleles. Controls are the same as those used in Fig. 3e.
**b**, Cells survival after HO pulse.

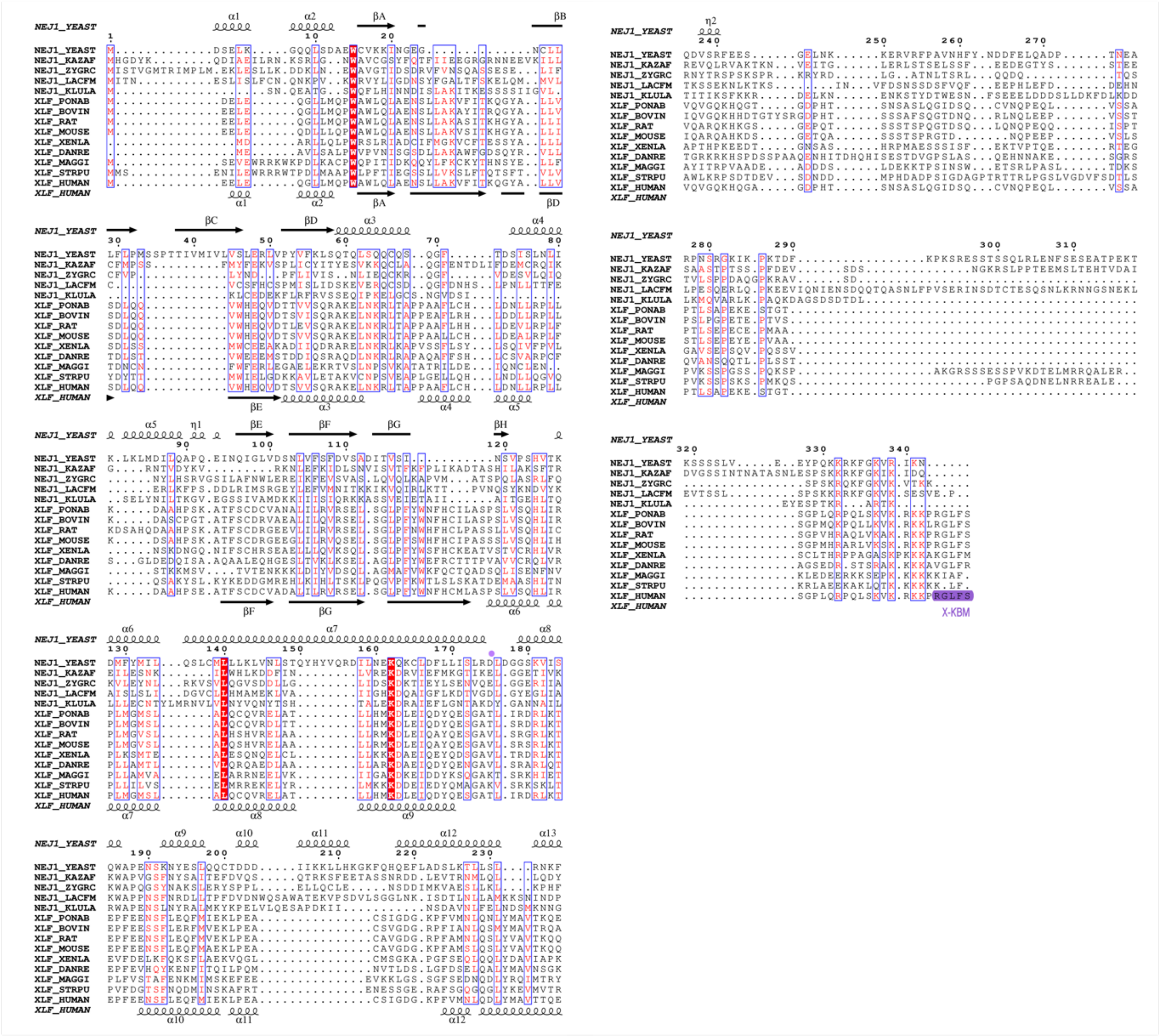


**Extended Data Fig. 7| Multiple sequence alignment of XLF/Nej1 homologues.**
The alignment was generated using the BLOSUM30 substitution matrix. The X-KBM motif in human XLF is highlighted in purple.

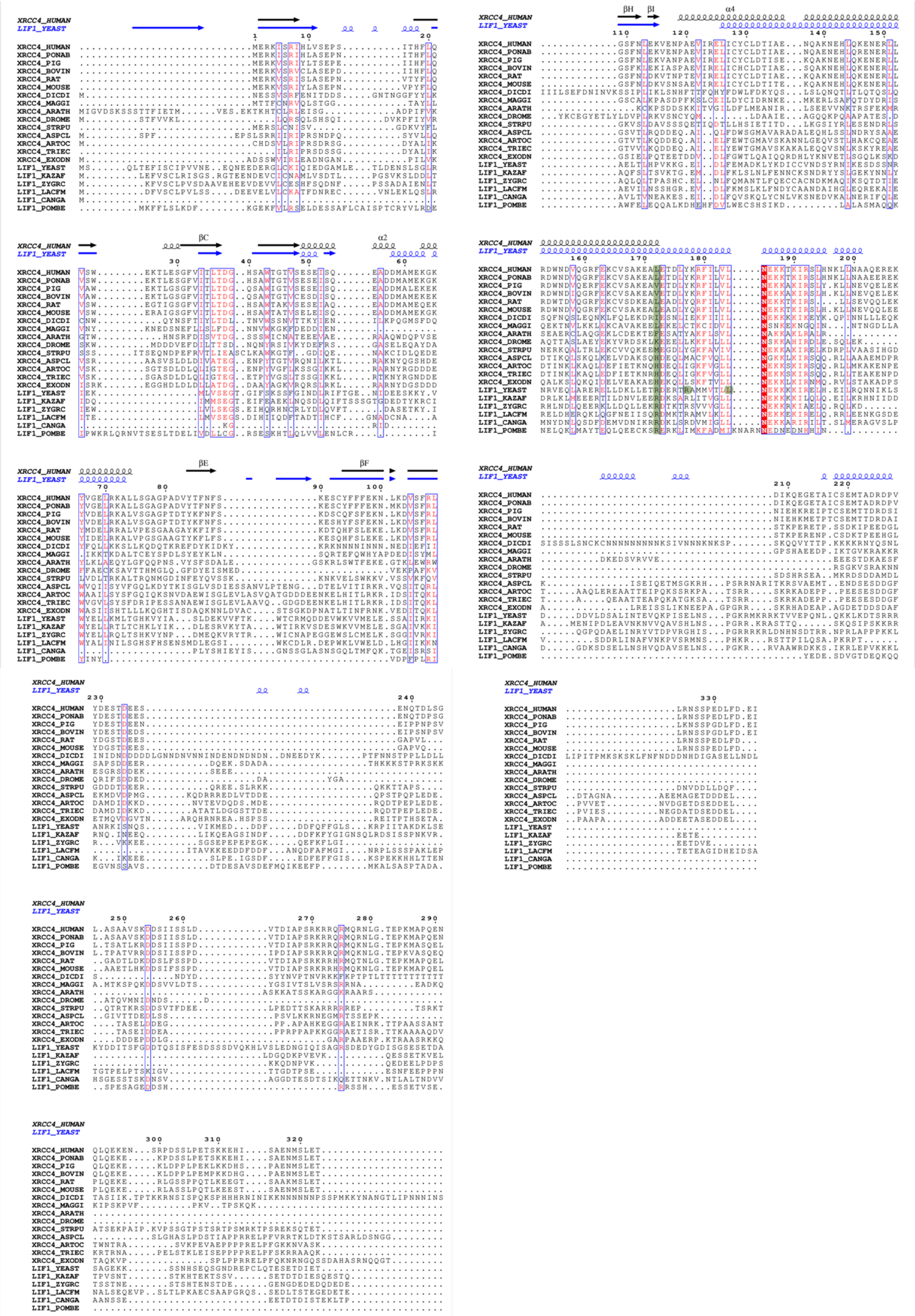

**Extended Data Fig. 8| Multiple sequence alignment of XRCC4/Lif1 homologues.**
The alignment was generated using the BLOSUM45 substitution matrix. Residue L173 in human XRCC4 (green) is conserved in higher eukaryotes and replaced by arginine or histidine in lower eukaryotes. Positions 177 and 181 were mutated to alanine for genetic studies and are highlighted in green in yeast.

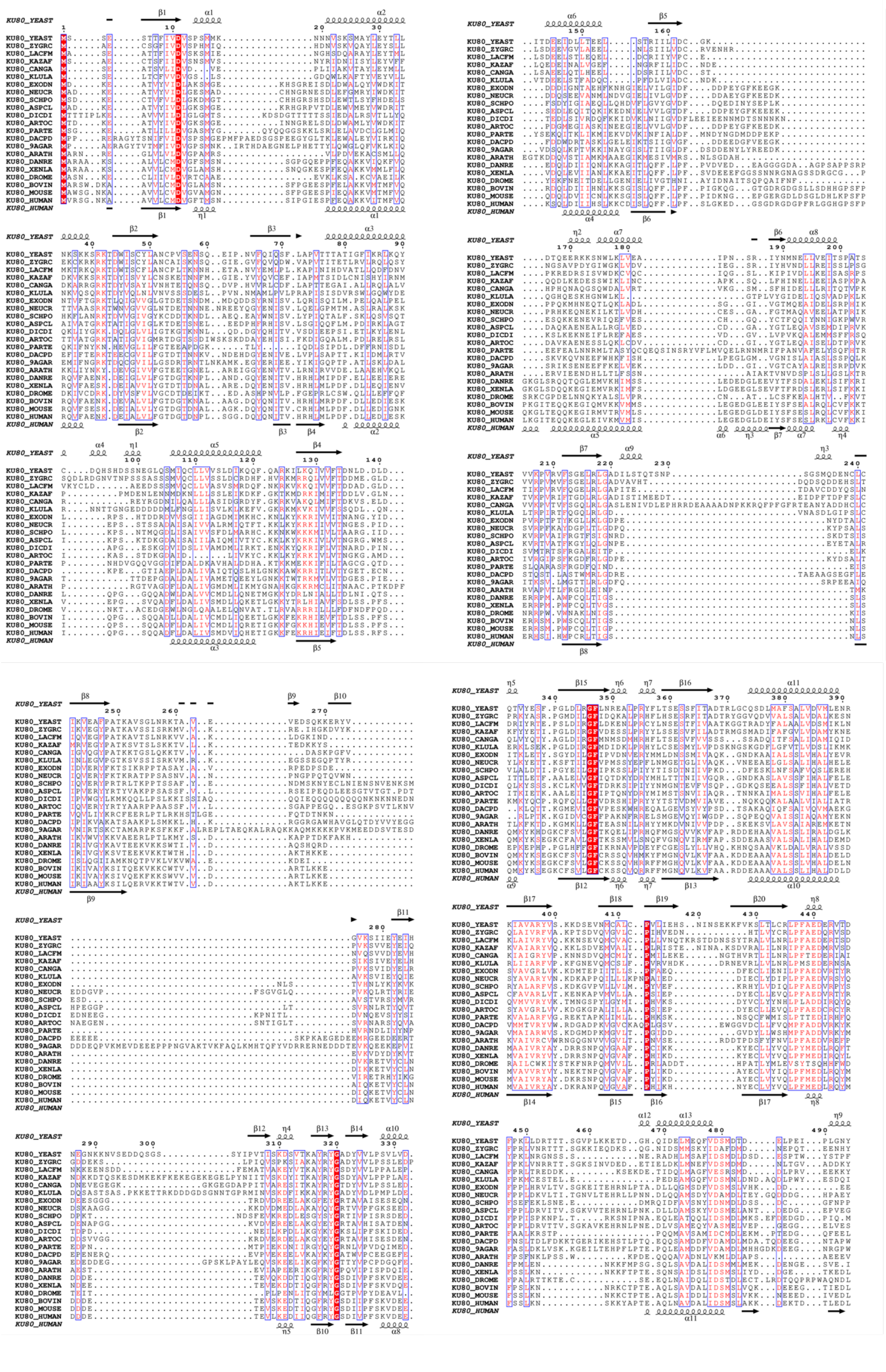

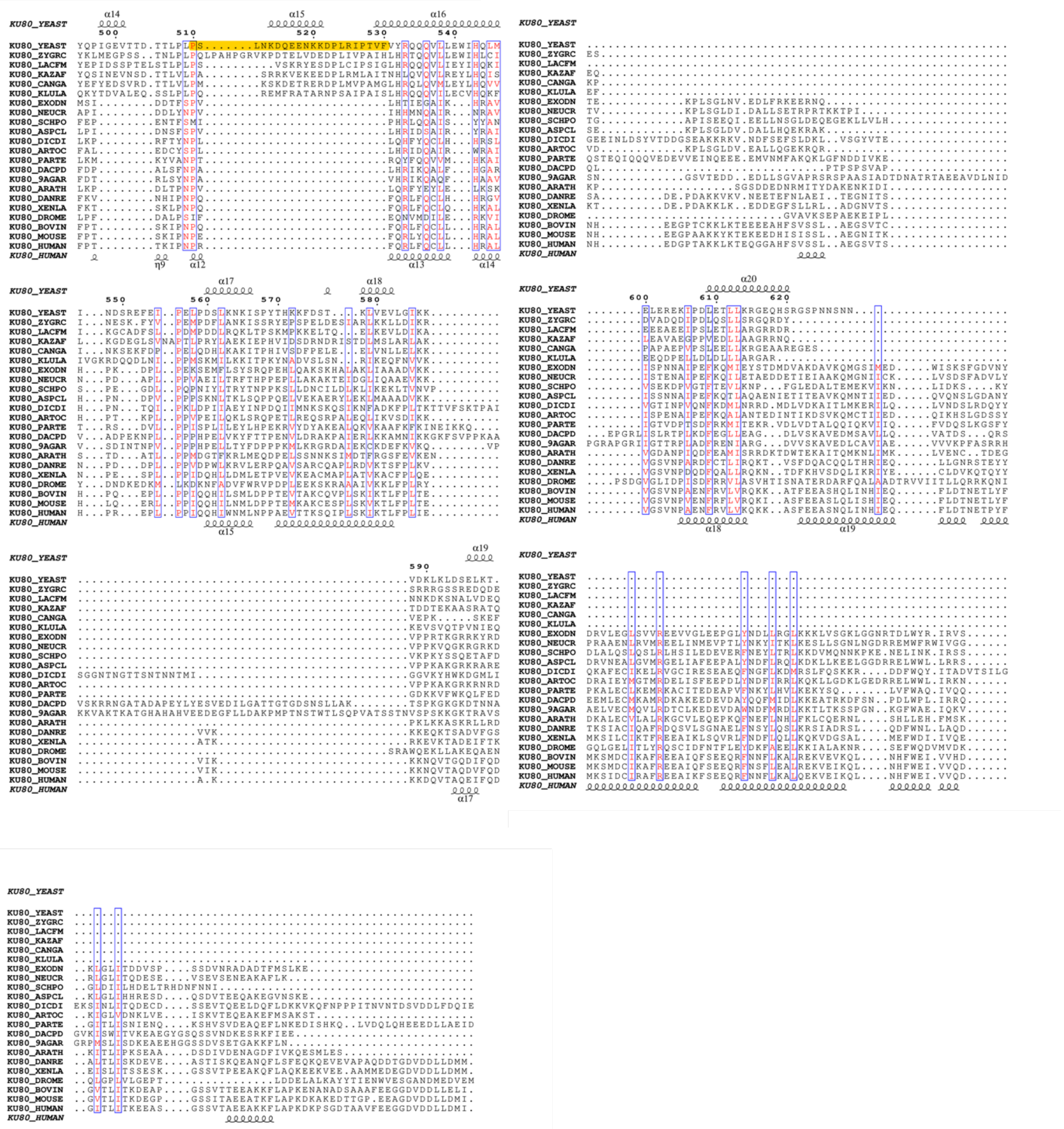

**Extended Data Fig. 9| Multiple sequence alignment of Ku80 homologues.**
The alignment was generated using the BLOSUM60 substitution matrix. Residues 510–530, corresponding to a loop present only in a subset of fungi, are highlighted in orange for *S. cerevisiae* Ku80 (yKu80).

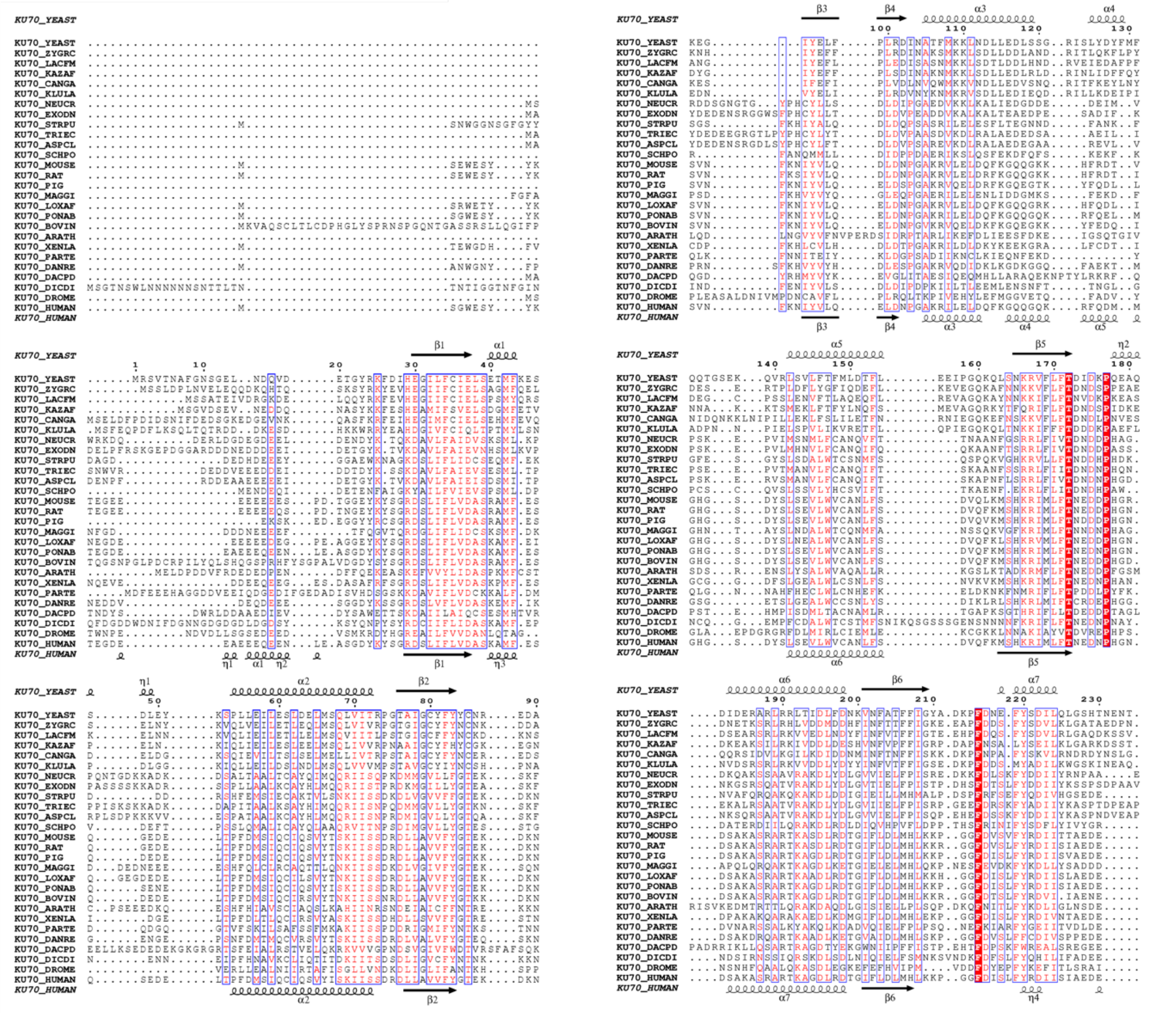

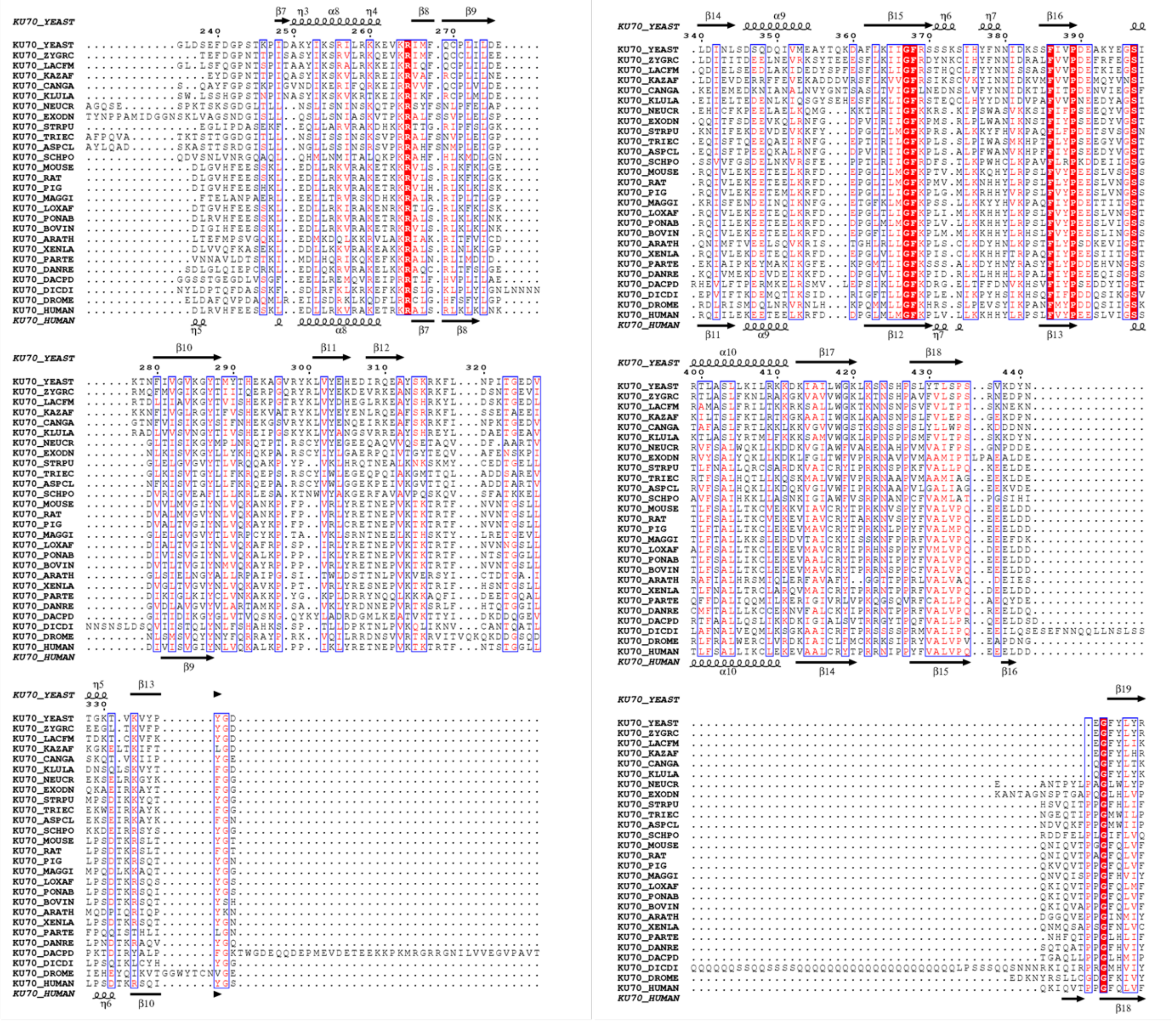

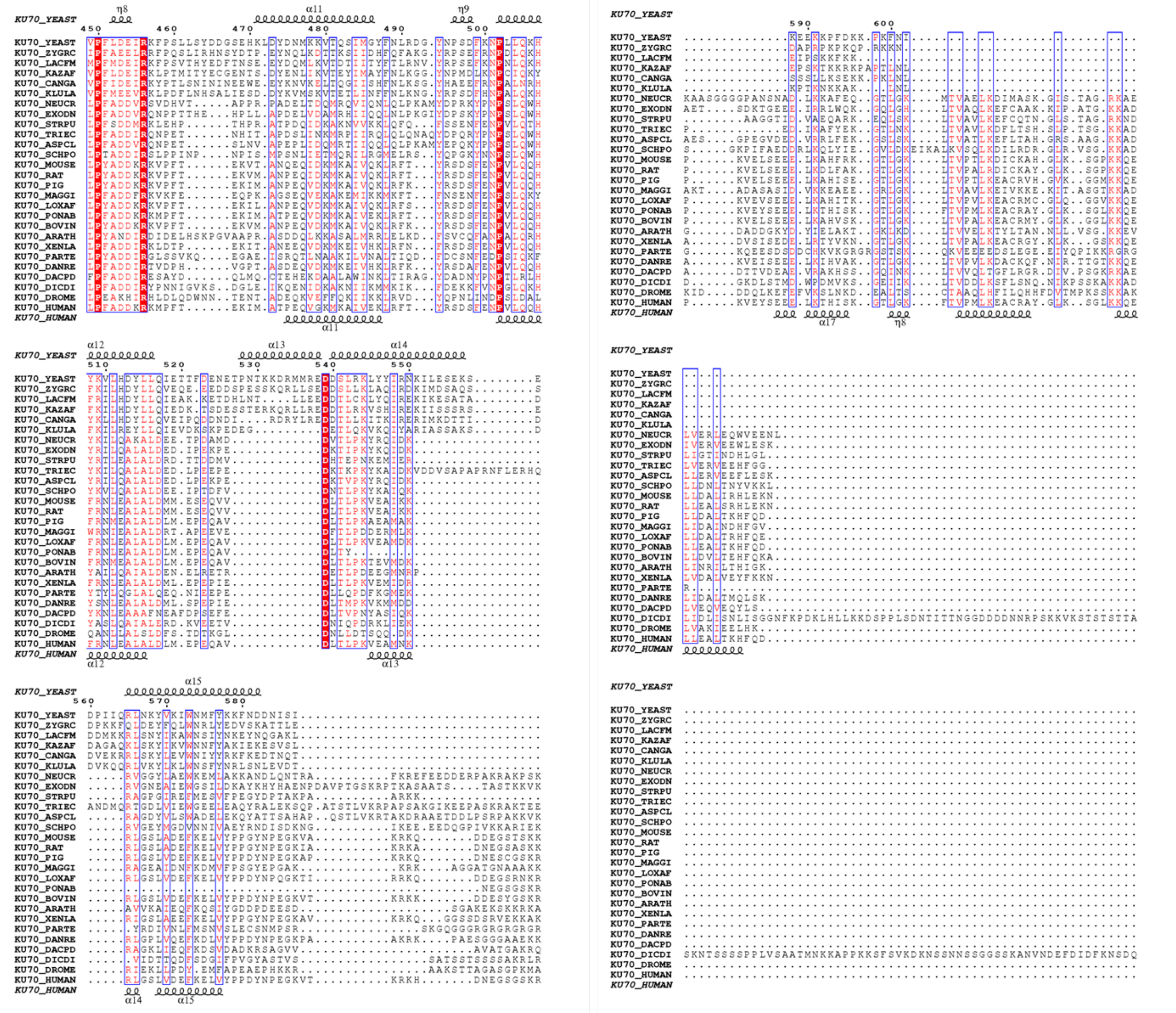

**Extended Data Fig. 10| Multiple sequence alignment of Ku70 homologues.**
The alignment was generated using the BLOSUM60 substitution matrix.

**Extended Data Table 1 Cryo-EM data collection, refinement and validation statistics**

| | LIGATION COMPETENT COMPLEX | DNA-ALIGNED PROTECTIVE COMPLEX 1 | DNA-ALIGNED PROTECTIVE COMPLEX 2 | DNA NON-ALIGNED PROTECTIVE COMPLEX |
|---|---|---|---|---|
| **Data collection and processing** | | | | |
| Microscope | Titan Krios G3i | | | |
| Detector | Falcon 4i | | | |
| Magnification | 130,000 | | | |
| Voltage (kV) | 300 | | | |
| Electron exposure ($e^{-}/Å^{2}$) | 50 | 40 | 40 | 40 |
| Defocus range (μm) | -0.4 to -2.4 | -0.4 to -2.4 | -0.4 to -2.4 | -0.4 to -2.4 |
| Pixel size (Å) | 0.96 | 0.96 | 0.96 | 0.96 |
| Initial particle images (no.) | 2,697,765 | 1,473,215 | 1,473,215 | 817,870 |
| Final particle images (no.) | 54,073 | 68,870 | 62,686 | 49,630 |
| Map resolution (Å) Mean<br>FSC threshold 0.143<br>$d_{99}$ | <br>3.34<br>3.0 | <br>3.34<br>3.2 | <br>3.32<br>3.2 | <br>5.77<br>4.6 |
| Map resolution range (Å) | 3.0 to 5.9 | 3.2 to 5.0 | 3.2 to 6.6 | 4.68 to 8.72 |
| **Refinement** | | | | |
| Initial model (PDB code) | 8S82, 1Z56 | 8S82,1Z56 | 8S82,1Z56 | 8S82,1Z56 |
| CC_mask | 0.70 | 0.68 | 0.72 | 0.56 |
| CC_box | 0.72 | 0.78 | 0.81 | 0.62 |
| CC_volume | 0.70 | 0.68 | 0.73 | 0.56 |
| CC_ligand | 0.70 | - | 0.51 | 0.40 |
| Model resolution (Å) FSC threshold 0.5 | 3.8 | 4.1 | 4.1 | 6.1 |
| Model composition<br>Non-H atoms<br>Protein residues<br>Nucleotides<br>Ligand (AMP) | <br>40897<br>4712<br>128<br>1 | <br>43437<br>5028<br>128<br>1 | <br>42980<br>4968<br>128<br>1 | <br>43876<br>5101<br>120<br>1 |
| *B* factors ($Å^{2}$)<br>Protein<br>Nucleotides<br>Ligand | <br>72.87<br>94.53<br>112.57 | <br>74.91<br>105.36<br>- | <br>84.93<br>98.93<br>134.85 | <br>83.54<br>98.92<br>131.40 |
| R.m.s. deviations<br>Bond lengths (Å)<br>Bond angles (°) | <br>0.004<br>1.048 | <br>0.005<br>1.140 | <br>0.006<br>1.155 | <br>0.005<br>1.057 |
| Validation<br>MolProbity score<br>Clashscore<br>Poor rotamers (%) | <br>1.96<br>3.00<br>4.23 | <br>2.00<br>3.92<br>3.97 | <br>2.05<br>3.69<br>4.94 | <br>1.70<br>4.35<br>0.04 |
| Ramachandran plot<br>Favored (%)<br>Allowed (%)<br>Disallowed (%) | <br>94.23<br>5.72<br>0.02 | <br>94.74<br>5.24<br>0.02 | <br>94.41<br>5.49<br>0.10 | <br>91.90<br>8.08<br>0.04 |

**Supplementary Table 1 Correlation coefficient between the model and the map, chain by chain**

CC plot of the ligation-competent complex

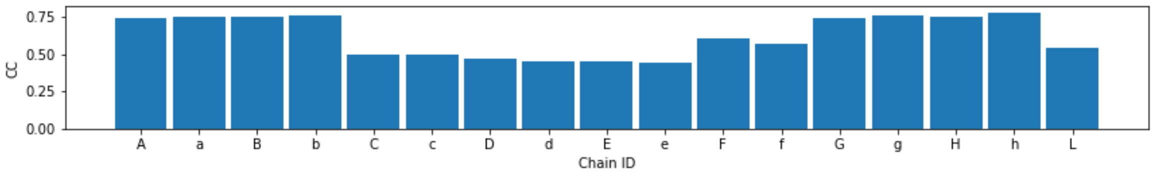


CC plot of the DNA-aligned protective complex 1

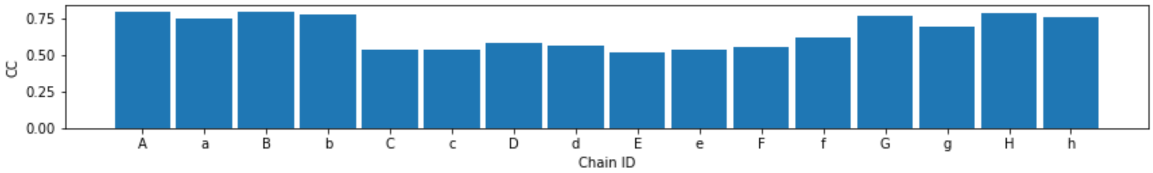


CC plot of the DNA-aligned protective complex 2

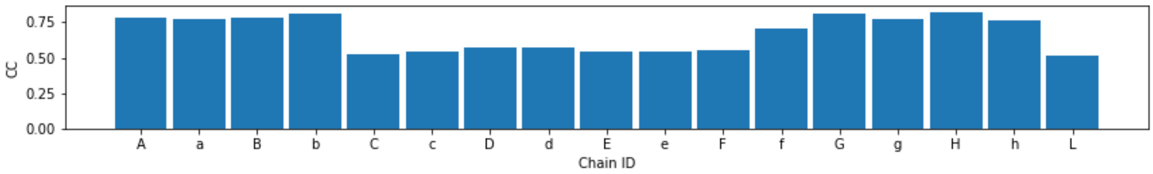


CC plot of the non-aligned protective complex

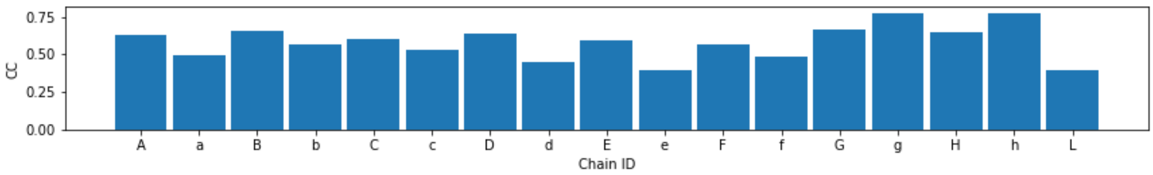


Chains
A&a: Ku80 B&b: Ku70
C&c, D&d: Lif1
E&e: Nej1
F&f: Lig4
G&g, H&h: DNA chains
L: the ligand AMP

**Supplementary Table 2**

DNA oligonucleotides list

| Name | Oligonucleotide sequences |
|---|---|
| **oSM0075** | 5'P-GTGTAATCTACTGACATCAGAGTTCTTAGATGCCCGG |
| **oSM0077** | CCGGGCAAGCTCGATCCCCGAGCTTCTAAGAACTCTGATGTCAGTAGATTACAddC |
| **oSM081'** | CCGGGCAAGCTCGATCCCCGAGCTTCTAAGAACTCTGATGTCAGTAGATTACACCCCddC |
| **oSM083** | CCGGGCAAGCTCGATCCCCGAGCTTCTAAGAACTCTGATGTCAGTAGATTACACGGGddG |
| **oSM089** | CCGGGCAAGCTCGATCCCCGAGCTTCTAAGAACTCTGATGTCAGTAGATTACACGGGG |

**Supplementary Table 3**

Strains list

| Name | Genotype |
|---|---|
| **Lev348** | *MATa ho∆ hml∆::ADE1 hmr∆::ADE1 ade1-100 leu2-3,112 lys5 trp1::hisG ura3-52 ade3::GAL::HO bar1∆ ::TRP1* |
| **Lev379** | *MATa ho∆ hml∆::ADE1 hmr∆::ADE1 ade1-100 leu2-3,112 lys5 trp1::hisG ura3-52 ade3::GAL::HO bar1∆ ::TRP1 lif1∆::KAN* |
| **Lev406** | *MATa ho∆ hml∆::ADE1 hmr∆::ADE1 ade1-100 leu2-3,112 lys5 trp1::hisG ura3-52 ade3::GAL::HO bar1∆ ::TRP1 nej1∆::KAN* |
| **Lev370** | MATa, ho∆, hml∆::ADE1 , hmr∆::ADE1, ade1-100,leu2-3,112 , lys5, trp1::hisG, ura3-52, ade3::GAL::HO, bar1∆::TRP1, yku80∆::KAN |

**Supplementary Table 4**

Plasmids list

| Name | Content | Origin |
|---|---|---|
| **PSC117** | *pRS315 pNEJ1_NEJ1-13Myc_tADH1* | this study |
| **PSC118** | *pRS315 pNEJ1_nej1-F71A-I114A-13Myc_tADH1* | this study |
| **PSC119** | *pRS315 pLIF1_LIF1-13Myc_tADH1* | this study |
| **PSC120** | *pRS315 pLIF1_lif1-F75A-I136A-13Myc_tADH1* | this study |
| **PSC164** | pRS315 yku80-Δloop (Δ510-523) + 13Myc | this study |
| **PSC165** | *pRS315* Yku80 + 13Myc | this study |
